# Cryo-EM structures and functional characterization of the lipid scramblase TMEM16F

**DOI:** 10.1101/455261

**Authors:** Carolina Alvadia, Novandy K. Lim, Vanessa Clerico Mosina, Gert T. Oostergetel, Raimund Dutzler, Cristina Paulino

**Author notes:** These authors contributed equally (Carolina Alvadia, Novandy K. Lim, Vanessa Clerico Mosina). Senior author (Raimund Dutzler, Cristina Paulino). Correspondence (R.D.), (C.P.).

## Abstract

The lipid scramblase TMEM16F initiates blood coagulation by catalyzing the exposure of phosphatidylserine in platelets. The protein is part of a family of membrane proteins, which encompasses calcium-activated channels for ions and lipids. Here, we reveal features of TMEM16F that underlie its function as lipid scramblase and ion channel. The cryo-EM structures of TMEM16F in Ca^2+^-bound and Ca^2+^-free states display a striking similarity to the scrambling-incompetent anion channel TMEM16A, yet with distinct differences in the catalytic site and in the conformational changes upon activation. In conjunction with functional data, we demonstrate the relationship between ion conduction and lipid scrambling. Although activated by a common mechanism, which likely resembles an equivalent process defined in the homologue nhTMEM16, both functions appear to be mediated by alternate protein conformations, which are at equilibrium in the ligand-bound state.

## INTRODUCTION

Lipid scramblases facilitate the movement of lipids between both leaflets of the bilayer (Bevers and Williamson, 2016; Nagata et al., 2016; Williamson, 2015), thereby dissipating the asymmetry present in the membrane. In contrast to ATP-dependent lipid flippases and floppases, scramblases are generally non-selective and do not require the input of energy. Lipid scrambling changes the properties of membranes and results in the exposure of negatively charged phospholipids to the outside, which are sensed by receptor proteins that in turn initiate important cellular responses (Nagata et al., 2016; Whitlock and Hartzell, 2016a). In platelets, scrambling is mediated by the protein TMEM16F, which catalyzes the externalization of phosphadidylserine (PS) in response to an increase of the intracellular Ca^2+^ concentration to trigger blood clotting (Bevers and Williamson, 2016; Suzuki et al., 2013). Splicing mutations of the TMEM16F gene were found to cause Scott syndrome, a severe bleeding disorder in humans and dogs (Brooks et al., 2015; Castoldi et al., 2011; Lhermusier et al., 2011). TMEM16F belongs to the TMEM16 protein family, whose members either function as anion-selective channels (Caputo et al., 2008; Schroeder et al., 2008; Yang et al., 2008) or lipid scramblases (Brunner et al., 2016; Falzone et al., 2018; Whitlock and Hartzell, 2016a). The structure of the fungal homologue nhTMEM16, determined by X-ray crystallography, has defined the general architecture of the family and provided insight into the mechanism of lipid translocation (Brunner et al., 2014). In nhTMEM16, each subunit of the homodimeric protein contains a membrane-accessible polar furrow termed the ‘subunit cavity’, which is believed to provide a suitable pathway for the polar lipid headgroups on their way across the hydrophobic core of the bilayer (Bethel and Grabe, 2016; Brunner et al., 2014; Jiang et al., 2017; Lee et al., 2018; Stansfeld et al., 2015; Pomorski and Menon, 2006). By contrast, single particle cryo-electron microscopy (cryo-EM) structures of TMEM16A, which instead of transporting lipids, solely facilitates selective anion permeation (Dang et al., 2017; Paulino et al., 2017a; Paulino et al., 2017b), revealed the structural differences that underlie the distinct function of this family branch. In TMEM16A, the rearrangement of an α-helix that lines one edge of the subunit cavity has sealed the membrane-accessible furrow, resulting in the formation of a protein-enclosed aqueous pore that is for a large part shielded from the bilayer. In both proteins, the binding of Ca^2+^ mediates activation of the catalytic site contained in each subunit, which in TMEM16A was shown to act as independent entity (Jeng et al., 2016; Lim et al., 2016). Whereas the structures of nhTMEM16 and TMEM16A have defined the architecture of the two most distant homologues of the family, TMEM16F appears, with respect to phylogenetic relationships, as intermediate between the two proteins. Although working as lipid scramblase (Suzuki et al., 2010; Watanabe et al., 2018), it is closer related to the ion channel TMEM16A than to nhTMEM16 (Figure S1A). Moreover, whereas scrambling-related ion conduction was found to be a feature of several family members (Lee et al., 2016; Malvezzi et al., 2018; Whitlock and Hartzell, 2016b), TMEM16F is the only lipid scramblase for which instantaneous calcium-activated currents were recorded in excised patches (Yang et al., 2012). To better understand how the small sequence differences in TMEM16F give rise to its distinct functional properties, we determined its structure by cryo-EM in Ca^2+^-bound and Ca^2+^-free states, both in a detergent and in a lipid environment. In parallel, we characterized the lipid transport properties of TMEM16F *in vitro* after reconstitution of the protein into liposomes, as well as ion conduction in transfected cells by electrophysiology. Collectively, our study reveals the architecture of TMEM16F, defines conformational changes upon ligand binding and suggests potential mechanisms for ion and lipid movement. In the most plausible scenario, both transport processes, although activated by the same mechanism, are mediated by distinct protein conformations which are at equilibrium in a calcium-bound state.

## RESULTS

### Functional properties of TMEM16F

In our study, we explored the relationship of the TMEM16F structure to its diverse functional properties. For this purpose, we have expressed TMEM16F in HEK293 cells and purified it in the detergent digitonin (Figures S1B and S1C). To confirm that the purified protein has retained its activity as a lipid scramblase, we have investigated lipid transport with proteoliposomes using an assay that was previously established for fungal TMEM16 scramblases (Malvezzi et al., 2013; Ploier and Menon, 2016). Our data reveals a Ca^2+^-induced activity, which, although dependent on the composition of proteoliposomes, facilitates transport of lipids with different headgroups, consistent with the broad lipid selectivity that was described for TMEM16F and other TMEM16 scramblases (Brunner et al., 2014; Malvezzi et al., 2013; Suzuki et al., 2013) (Figures S1D, S1E and S1F). Activation by Ca^2+^ is instantaneous and saturates with an EC_50_ of about 2 µM (Figures 1A, 1B and S1G). Thus, our experiments demonstrate the function of purified and reconstituted TMEM16F as a Ca^2+^-activated lipid scramblase.

**Figure 1.**
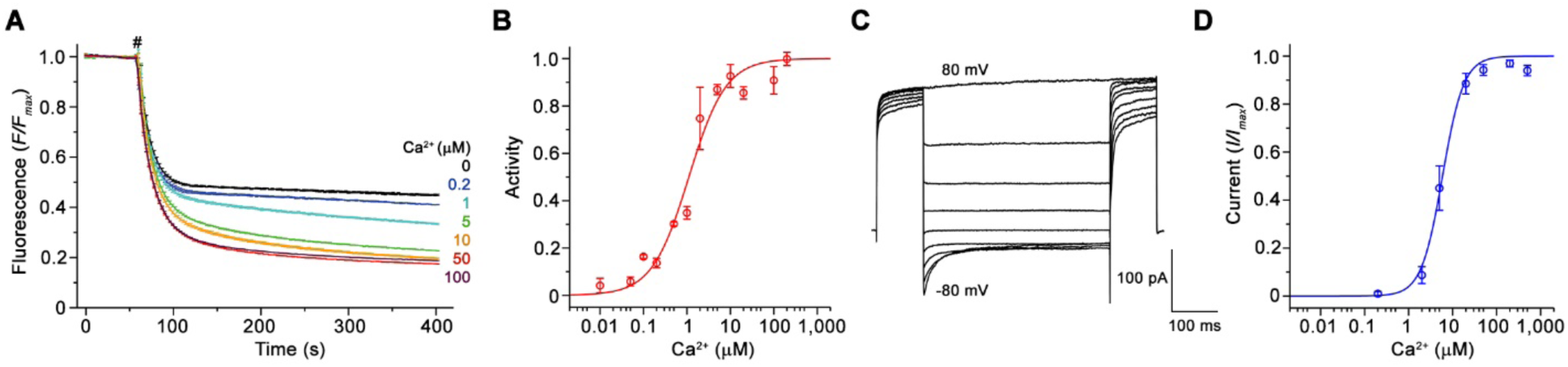
Functional characterization of TMEM16F. (A) Ca^2+^-dependence of scrambling activity in TMEM16F-containing proteoliposomes. Traces depict fluorescence decrease of tail-labeled NBD-PE lipids after addition of dithionite (#) at different Ca^2+^ concentrations. Data show averages of three technical replicates. (B) Ca^2+^ concentration-response relationship of TMEM16F scrambling. Data show mean of six independent experiments from three protein reconstitutions. (C) Representative current traces of TMEM16F at 200 µM Ca^2+^ recorded from inside-out patches. (D) Ca^2+^ concentration-response relationship of currents. Data show mean of seven biological replicates. A, B, D, errors are s.e.m.. See also Figure S1.

Besides its capability to transport lipids, TMEM16F was also reported to act as an ion channel (Kunzelmann et al., 2014; Scudieri et al., 2015; Yang et al., 2012; Yu et al., 2015). To recapitulate these properties, we have studied ion conduction in excised patches in the inside-out configuration. As previously described, TMEM16F mediates currents that are activated in response to elevated intracellular Ca^2+^ concentrations (Yang et al., 2012) (Figures 1C, 1D and S1H). Similar to scrambling, the activation is instantaneous and currents saturate with an EC_50_ of 8 µM and a Hill coefficient of about 2 (measured at 80 mV, Figure 1D). Unlike for the anion selective TMEM16A, the currents are slightly selective for cations and they retain their strong outward rectification in the entire ligand-concentration range (Figures 1C and S1I). Whereas the mild rectification of instantaneous currents is likely a consequence of conduction through an asymmetric pore (Paulino et al., 2017b), the presence of time-dependent relaxations in response to changes of the transmembrane voltage and the increased rectification at steady-state reflect a voltage-dependent transition that is not obviously linked to ligand binding (Figure 1C). As the activation of ion conduction and lipid scrambling occurs in a similar ligand concentration range they both likely rely on the same mechanism of Ca^2+^-activation.

### TMEM16F structure

Since TMEM16F combines the functional properties of a lipid scramblase and an ion channel, we were interested in the structural features that distinguishes it from the anion-selective channel TMEM16A and the fungal scramblase nhTMEM16. For that purpose, we have determined structures of TMEM16F in the absence and presence of ligand (Table S1). Structures of the protein in the detergent digitonin were determined for Ca^2+^-bound and Ca^2+^-free states by cryo-EM at 3.2 Å and 3.6 Å, respectively (Figures 2, S2, S3, Table S1 and Movie S1 and S2). Both structures are very similar, except for conformational differences in the ligand-binding site and the presumed catalytic region (Figure 2C). To investigate the effect of a lipid bilayer on the structural properties of TMEM16F, we reconstituted the protein into large 2N2 lipid nanodiscs containing a lipid composition that retains scrambling activity, although with slower kinetics, (Figure S1E) and collected cryo-EM data of both states in a membrane-like environment (Figures S4 and S5).

**Figure 2.**
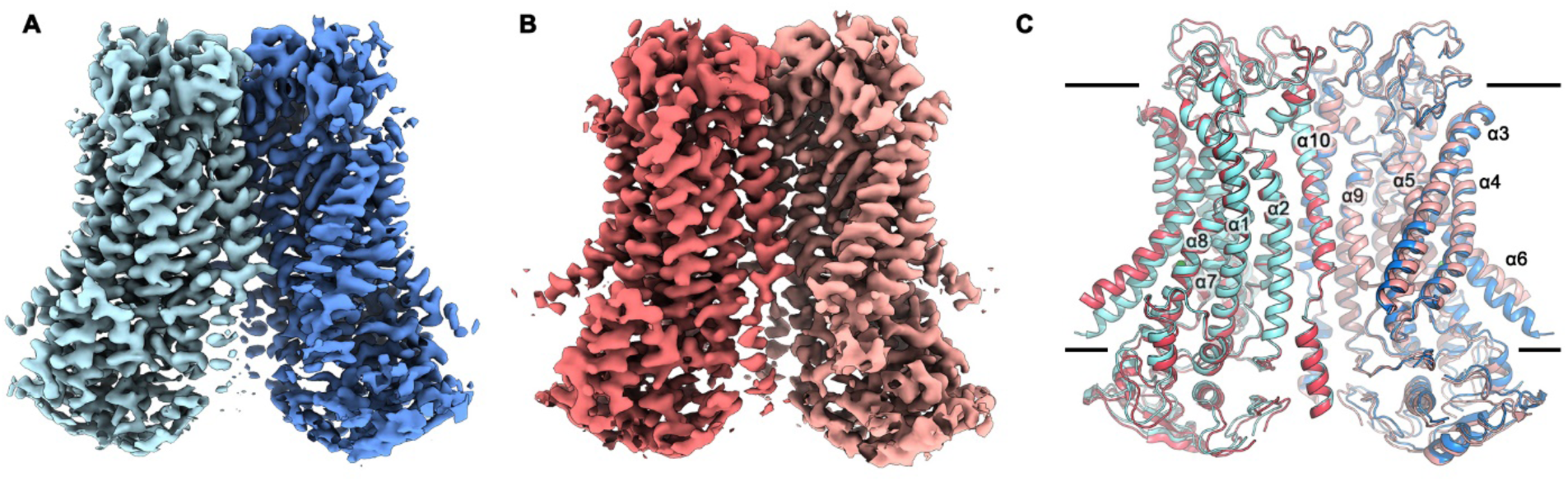
TMEM16F structures. (A and B) Cryo-EM map of TMEM16F in digitonin in presence (A) and absence (B) of calcium at 3.2 Å and 3.6 Å, respectively. (C) Cartoon representation of a superposition of the Ca^2+^-bound (blue) and Ca^2+^-free TMEM16F (red) dimer. Transmembrane helices are labelled and the membrane boundary is indicated. See also Figures S2 to S6, Table S1 and Movie S1 and S2.

Whereas nominally of similar resolution as our structures determined in detergents, the preferred orientation of the protein in nanodiscs on the grids resulted in anisotropic cryo-EM maps, in which some of the details observed in the digitonin samples are blurred (Figures S4-S6). This problem is less pronounced in the nanodisc data obtained in presence of Ca^2+^ than in the dataset of the Ca^2+^-free sample. Nevertheless, the data in nanodiscs are of sufficient quality to allow for a comparison of corresponding states of TMEM16F in a detergent and lipid environment. For most parts, these structures are virtually indistinguishable except for α-helices 3 and 4, which show a larger tilt towards α-helix 6 in the cryo-EM maps of nanodisc samples, a feature that is more prominent in the Ca^2+^-free state (Figure S6C). While our datasets reveal an effect of lipids on parts of the structure, the overall similarity demonstrates a general equivalence of the protein conformation in detergent and in a membrane environment (Figure S6A and S6C). In addition, we were interested whether we could identify any structural flexibility within each of the four datasets. While no heterogeneity was found in the data in absence of Ca^2+^ (Figures S3I and S5I), we were able to identify distinct conformations for α-helices 3 and 4 in presence of Ca^2+^ (Figures S2I and S4I). These differences are more pronounced in the nanodisc sample and could point towards a structural transition into a more open conformation of the ‘subunit cavity’, as observed in the scramblase nhTMEM16 (see accompanying manuscript), although such conformation is not found in our data (Figure S6C). In contrast to TMEM16A (Paulino et al., 2017b) and nhTMEM16 (see accompanying manuscript), no obvious distortion of either the micelle or lipid nanodiscs was observed in the maps of TMEM16F (Figures S2H, S3H, S4H and S5H). However, in light of the structural similarity, a destabilization of the membrane by the protein, as described for nhTMEM16, is plausible.

With a pairwise sequence identity of 38%, the relationship of TMEM16F is closer to the anion channel TMEM16A (Figure S1A), than to the more distantly related fungal scramblase nhTMEM16 and its mammalian orthologue TMEM16K, with which it shares sequence identities of 21% and 25%, respectively. It may thus not be surprising that, with respect to its general architecture, TMEM16F closely resembles TMEM16A. The similarity encompasses all parts of the protein and is particularly pronounced in the Ca^2+^-bound conformation, where TMEM16F superimposes with root mean square deviations (RMSDs) of 2.2 Å and 4.0 Å on TMEM16A and nhTMEM16, respectively (Figure 3). The structural features in the ligand-bound state, which distinguish the membrane-accessible furrow in nhTMEM16 and the protein-enclosed pore in TMEM16A, are believed to be the hallmark that differentiate lipid scramblases from ion channels in the family (Paulino et al., 2017b). It is thus remarkable to find equivalent helices in TMEM16F in a similar conformation as in TMEM16A (Figure 3B and 3C). In light of their strong structural similarity, the question is thus pertinent why TMEM16F but not TMEM16A catalyzes lipid movement. It also remains uncertain, whether the here captured Ca^2+^-bound structure of TMEM16F might show a not fully activated state, or whether the scrambling mechanism of TMEM16F may differ from nhTMEM16 and its human orthologues.

**Figure 3.**
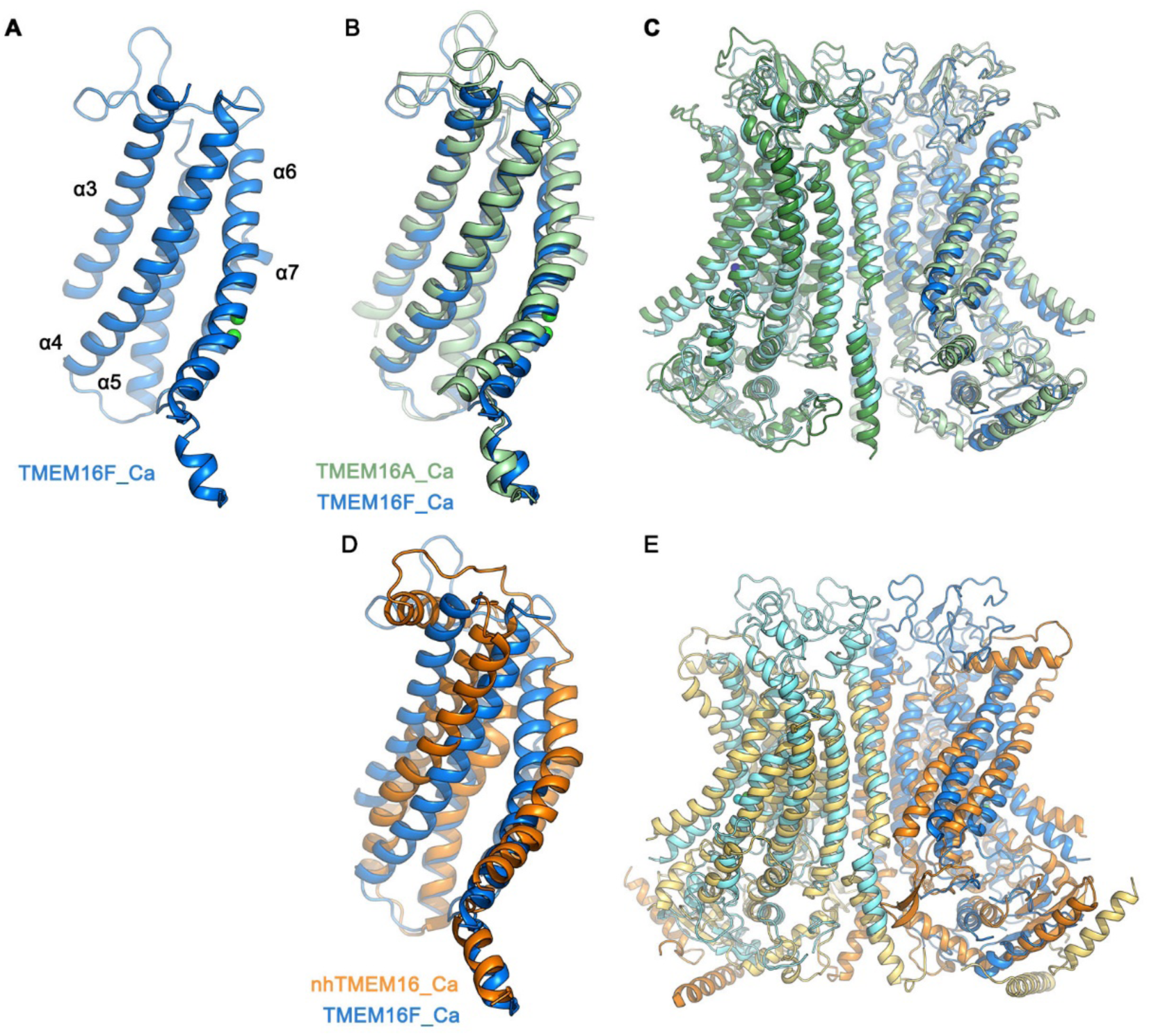
Structural comparison of TMEM16 homologues. (A) Subunit cavity in the Ca^2+^-bound TMEM16F structure composed of α-helices 3–7 (blue). (B and C) Superposition of the Ca^2+^-bound structures of TMEM16F (blue) and the anion channel TMEM16A (PDBID 5OYB (Paulino et al., 2017a), green). (B) ‘subunit cavity’, (C) dimeric protein. (D and E) Superposition of the Ca^2+^-bound structures of TMEM16F (blue) and the fungal scramblase nhTMEM16 (PDBID 4WIS (Brunner et al., 2014), orange). (D) ‘subunit cavity’, (E) dimeric protein.

### Architecture of the catalytic region

Although the relation of the Ca^2+^-bound structure of TMEM16F to a defined functional state of the protein is at this stage ambiguous, its comparison to the structures of family members obtained at equivalent conditions reveals interesting parallels. The architecture of the putative catalytic unit of TMEM16F in a Ca^2+^-bound conformation shows a strong structural resemblance to the ion permeation path in TMEM16A (Figure 3B). In TMEM16A, the hourglass-shaped pore contains two aqueous vestibules on either side of the membrane that are bridged by a narrow neck (Paulino et al., 2017a). A related architecture is observed in TMEM16F, where closer interactions between α-helices and the presence of bulky residues along the path further constrict the pore (Figures 4A to 4F). Thus, it is still unclear whether the observed conformation of TMEM16F could facilitate ion conduction. In TMEM16A, a strong positive electrostatic potential throughout the pore, originating from an excess of cationic residues, determines its selectivity for anions (Paulino et al., 2017a). Due to a changed distribution of charges in the pore region of TMEM16F, the potential is generally less positive, which is consistent with the poor selectivity of currents passed by this protein (Figures 4G and S1I).

**Figure 4.**
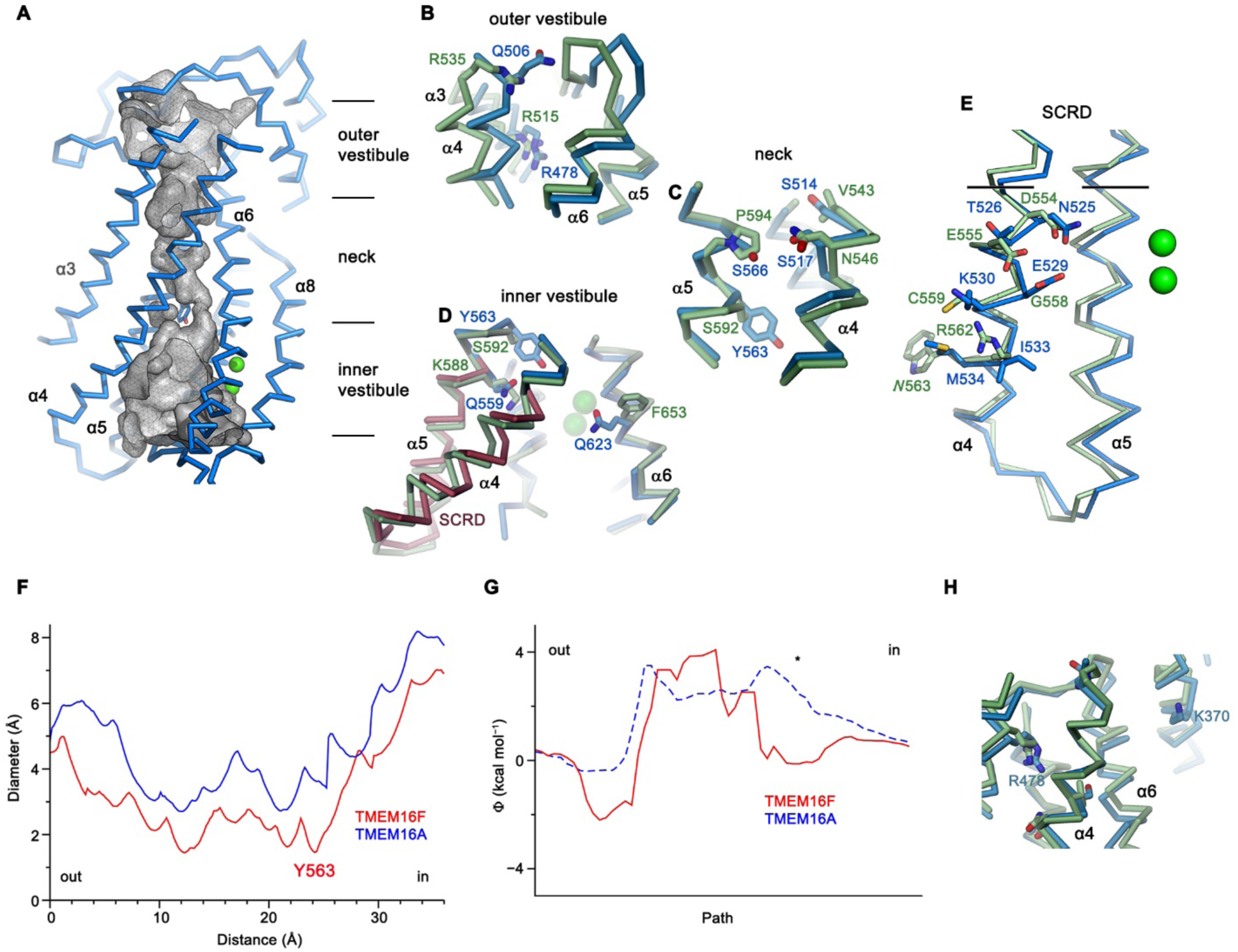
Catalytic site. (A) Putative ion conduction pore in TMEM16F. Grey mesh shows the surface of the pore (sampled with probe radius of 1 Å). (B to D) Superposition of selected pore regions of TMEM16F (blue) and the anion the anion channel TMEM16A (green, remodeled version of PDBID 5OYB see Methods) along the extracellular vestibule (B), the neck (C) and the intracellular vestibule with the 35 amino acid long scrambling domain (SCRD) on α-helix 4-5 of TMEM16F highlighted in brown (D). (E), View of the SCRD rotated by about 90 degrees relative to (D). The SCRD boundary is indicated by black lines. (E) Diameter of the putative pores of TMEM16F (red) and TMEM16A (blue) as calculated by HOLE (Smart et al., 1996) from the extracellular (out) to the intracellular side (in). The position of Tyr 563 is indicated. (F) Electrostatic profile calculated along the pore regions of TMEM16F (red) and TMEM16A (blue, dashed). The potential is less positive in TMEM16F for most of the pore, consistent with the observed slight preference for cations over anions. * indicates location of the Ca^2+^-binding site. (H) Location of Lys 370 in relation to the pore region of TMEM16F (blue) in a superposition with TMEM16A (green). (A to E and H), Selected residues are displayed as sticks, bound calcium ions as green spheres.

Whereas in the extracellular vestibule, the uncharged Gln 506, which has replaced an arginine at the equivalent position in TMEM16A, lowers the positive charge density in the outer mouth of the pore, Arg 478 at the extracellular boundary to the neck is conserved (Figure 4B). The neck of TMEM16F shares a similar amphiphilic character as in TMEM16A but it contains several changes, which decrease its hydrophobicity (Figure 4C). Further towards the cytoplasm, the bulky Tyr 563 lowers the pore diameter of TMEM16F to 1.6 Å (Figures 4A, 4D and 4F), which would have to expand to permit ion conduction. Finally, at the boundary to the wide intracellular vestibule, Gln 559 replaces a lysine at the equivalent position in TMEM16A, thereby removing another positive charge from the putative ion permeation path (Figure 4D).

Similar to its paralogue TMEM16A, the interacting α-helices 4 and 6 disengage towards the cytoplasm and open a hydrophilic gap to the membrane, which causes a dilation of the pore at the intracellular vestibule (Figures 4D and 4F). In TMEM16F this region contains the scrambling domain (SCRD), encompassing the intracellular parts of α-helices 4 and 5, which was previously described as determinant for lipid movement (Gyobu et al., 2017; Yu et al., 2015) (Figures 4D and 4E). The intracellular gap of TMEM16F also parallels the equivalent region of nhTMEM16 (Figures 3D and 3E), but the closure of the cavity towards the extracellular side interrupts the polar and membrane-accessible furrow found in the Ca^2+^-bound ‘open state’ of the fungal scramblase and instead resembles its Ca^2+^-bound intermediate observed in nanodiscs (see accompanying manuscript). The region of the closed cavity that faces the lipid bilayer in TMEM16F is hydrophobic and resembles TMEM16A, except for few residues such as Gln 623, located on α-helix 6 facing the SCRD, and Lys 370 located at the extracellular domain close to the membrane boundary (Figures 4D and 4H).

### Calcium activation

The structure of TMEM16F in the presence of ligand defines the location of two Ca^2+^ ions per subunit that are bound to a site, which strongly resembles its counterpart in nhTMEM16 (Brunner et al., 2014) and TMEM16A (Paulino et al., 2017a) (Figures 5 and S6B). In this site, conserved acidic and polar residues located on α-helices 6–8 coordinate the ions with similar geometry. Different from its paralogue functioning as anion channel, TMEM16F does not contain an insertion of a residue in α6 near the hinge (Figure S1A), which causes a partial unwinding and the formation of a π-bulge in TMEM16A (Paulino et al., 2017a). Consequently, the helix in the ligand-bound conformation of TMEM16F might be in a less strained conformation.

**Figure 5.**
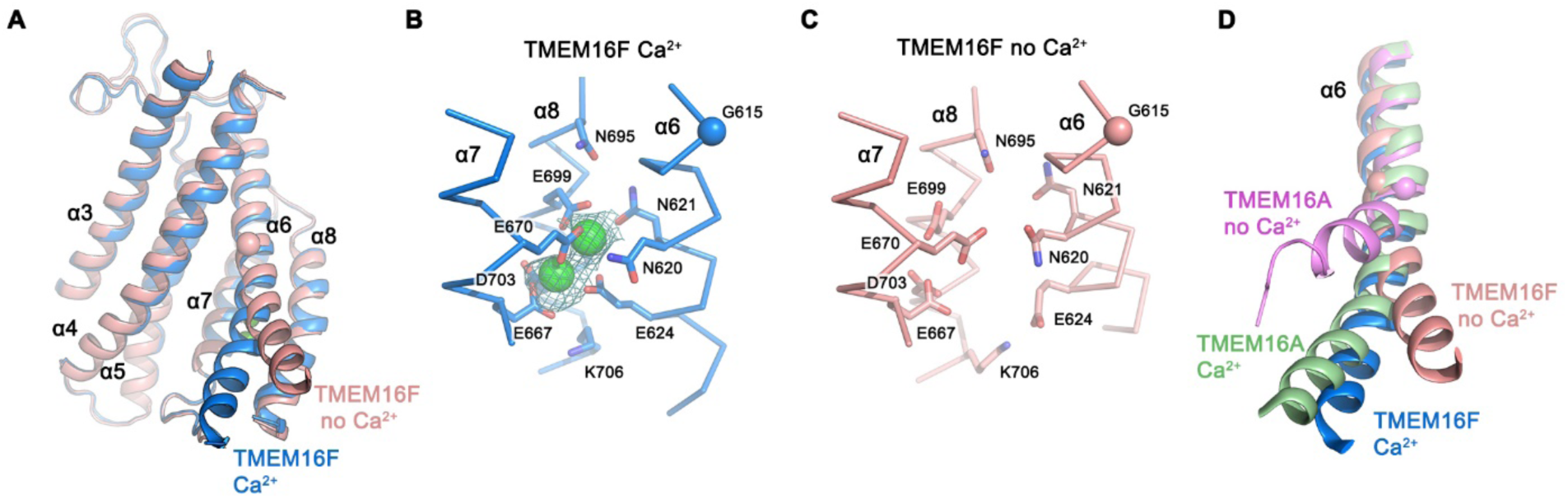
Ca^2+^-binding site and conformational changes. (A) Superposition of α-helices 3–8 of TMEM16F in the presence (blue) and absence of Ca^2+^ (coral). (B and C) Close-up of the Ca^2+^-binding site in complex (B) or in absence of ligand (C). Coordinating residues are depicted as sticks, calcium ions as green spheres (with surrounding cryoEM density contoured at 5σ). (D) Comparison of Ca^2+^-induced movements in TMEM16F (Ca^2+^-bound in blue, Ca^2+^-free in coral) and TMEM16A (Ca^2+^-bound (PDBID 5OYB) in green, Ca^2+^-free (PDBID 5OYG) in pink). Shown is a cartoon superposition of α-helix 6. (A to D) Pivot for movement around Gly 615 is depicted as sphere.

The conformational changes that take place in TMEM16F upon ligand binding can be appreciated in the comparison of Ca^2+^-bound and Ca^2+^-free conformations (Figures 2C, 5A-5C, and S6). As in TMEM16A, the absence of the ligand causes a local transition that is confined to the pore region, whereas the remainder of the structure remains largely unaffected (Figs. 2C, 5A and S6C). Besides comparably small rearrangements of α3 and α4, which are less pronounced in the nanodisc structures (Figure S6C), the largest difference is found at the intracellular half of α6 (Figure 5A). In the presence of Ca^2+^, the bound ligands provide an interaction platform for negatively charged and polar residues on α6, thereby immobilizing the helix in the observed conformation (Figure 5B). In the ligand-free state, this helix has altered its conformation in response to the loss of the positively charged interaction partner, leading to an increase in mobility as indicated by the weaker density (Figure S6). The transition of α6 can be described by a rigid-body movement of its intracellular half by 20°, around a hinge located close to a conserved glycine residue (Gly 615, Figures 5 and S1A), whose equivalent in TMEM16A was shown to be critical for conformational rearrangements (Paulino et al., 2017a). However, in contrast to TMEM16A, the observed change in TMEM16F is considerably smaller, proceeds in the opposite direction and it is not accompanied by a rotation around the helix axis (Figure 5D). Thus, whereas in the apo-form of TMEM16A α6 approaches α4, thereby closing the gap between both helices, the movement in TMEM16F is directed away from α4 (Figures 5A and 5D). The observed structural transition opens the access of the ion binding site to the cytoplasm and widens the gap to the SCRD, which likely contributes to the inhibition of scrambling by increasing the barrier for lipids by a currently unclear mechanism. The general structural resemblance of the Ca^2+^-free conformation of TMEM16F to the equivalent structure of nhTMEM16 determined in lipid nanodiscs (see accompanying manuscript) underlines their correspondence to an inactive state of the respective protein.

### Functional characterization of mutants affecting activation, ion conduction and scrambling

Whereas the described structures have revealed the detailed architecture of TMEM16F and its conformational changes in response to ligand binding, the similarity of the protein to its paralogue working as anion channel leaves important aspects of structure-function relationships ambiguous.

The first question we have addressed was whether Ca^2+^-binding to the conserved site of TMEM16F would affect scrambling and conduction in a similar manner. We have thus investigated the scrambling activity of mutants of the binding site and found a strong decrease in the potency of Ca^2+^ in the mutant E667Q located on α7 and little detectable activity in the mutant E624Q on α6 (Figures 5B, 6A, S7A and S7B), consistent with a very similar phenotype of equivalent mutants on the activation of the chloride channel TMEM16A (Brunner et al., 2014; Lim et al., 2016; Tien et al., 2014; Yu et al., 2012). By using patch-clamp electrophysiology, we found a comparable effect of the same mutations on the activation of currents, in line with previous observations (Yang et al., 2012), thus emphasizing the mutuality in the regulation of both processes (Figures 5B, 6B, S7C and S7D). We next investigated the role of the observed conformational change of α6 on activation and the importance of the conserved glycine position as hinge for its movement. In patch-clamp experiments, the mutation of the conserved Gly 615 to alanine increased the potency of Ca^2+^ (Figures 6B and S7E), indicating a stabilization of the open state upon mutation of this flexible position, which parallels a behavior observed in the channel TMEM16A (Paulino et al., 2017a). Hence, the effect of mutations on the Ca^2+^-binding site and the gating hinge indicate that the conformational changes observed in α6 are the basis of a common mechanism of regulation for scrambling and ion conduction in TMEM16F.

**Figure 6.**
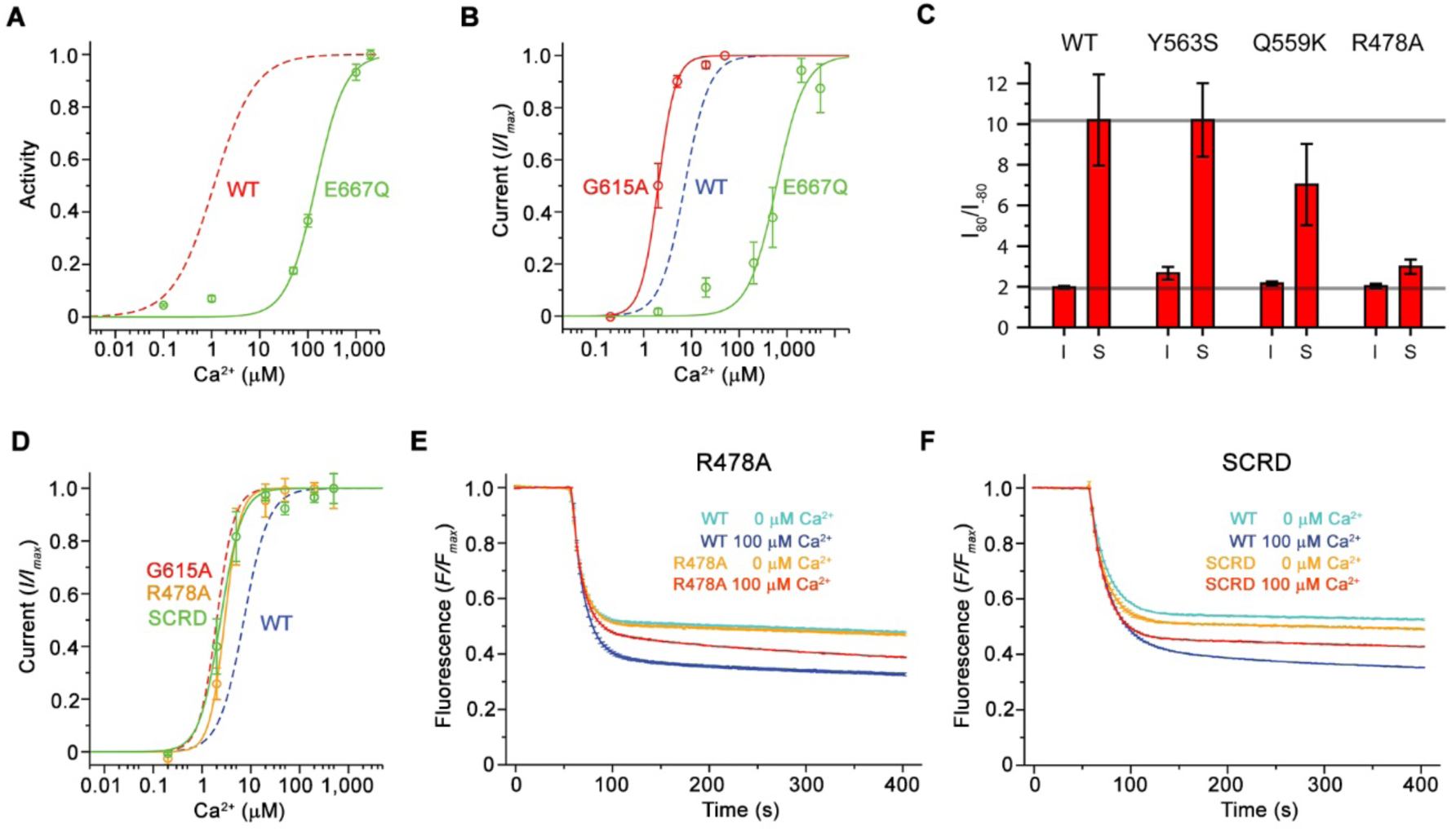
Functional properties of mutants. (A) Ca^2+^ concentration-response relationship of lipid scrambling in the binding site mutant E667Q. Data show average of either six (0, 1, 10, 100, 1000 µM) or three (other concentrations) independent experiments from two protein reconstitutions. (B) Ca^2+^ concentration-response relationship of currents of E667Q (n=4) and the hinge residue G615A (n=4). (A and B) For comparison WT traces are shown as dashed lines. (C) Rectification indices (I_80m_V/I–_80mV_) of instantaneous (I) and steady-state currents (S) of WT (n=12) and the mutants Y563S (n=8), Q559K (n=10) and R478A (n=10). (D) Ca^2+^ concentration-response relationship of currents of mutants R478A (n=5) and TMEM16^FSCRD^ (SCRD, n=6). For comparison WT and G615 traces are shown as dashed lines. (E and F) Scrambling of the mutants R478A (E) and TMEM16F^SCRD^ (SCRD) (F). (E and F) Data show average of three independent experiments of one protein reconstitution. WT reconstituted in a separate set of liposomes from the same batch is shown in comparison. (B and D) Data show averages from independent biological replicates. (A to F), Errors are s.e.m.. See also Figure S7.

Further, we were interested in the structural elements mediating both transport processes. The resemblance of the pore region of TMEM16F to TMEM16A, suggests that ion conduction could proceed at the same location. We have thus investigated the influence of mutations of pore-lining residues on ion conduction, by either truncation to alanine or replacement with their equivalents in TMEM16A. While Tyr 563 constricts the pore diameter of TMEM16F (Figures 4A, 4C and 4F), the mutation Y563S did not alter the rectification properties of currents in a noticeable manner (Figures 6C and S7F). The mutation of Gln 559 to lysine at the boundary between the intracellular vestibule and the narrow neck (Figure 4D), influences the selectivity of TMEM16F (Figure S7G), as described previously (Yang et al., 2012). In the mutant Q559K, we find a moderate decrease in the rectification of steady-state currents, an effect that is particularly pronounced in a subset of the data (Figures 6C and S7F). Finally, when mutating the conserved Arg 478 (Figure 4B) at the extracellular entrance of the narrow neck to alanine, the potency of Ca^2+^ is increased and voltage-dependent relaxations, and consequently the strong rectification at steady-state, largely disappears (Figure 6C, 6D, S7F and S7H). The effect of the mutations of Gln 559 and Arg 478 on the steady state rectification is consistent with the participation of these residues in structural transitions of the protein, which influence the open probability of the ion conduction pore in a voltage-dependent manner.

As the mutation R478A at the extracellular part of the pore affected channel properties, we next investigated its effect on lipid scrambling. Strikingly, the mutant strongly decreased the kinetics of scrambling (Figures 6E and S7I), consistent with previous data obtained from cellular assays (Gyobu et al., 2017). In contrast to the pronounced effect of a charged residue within the pore region of TMEM16F, the mutation of Lys 370, which is located in proximity to α6 at the extracellular end of the putative lipid translocation path outside of the pore, did not show a detectable effect on scrambling (Figures 4H, S7I and S7J).

Consistent with previous data from cellular assays (Yu et al., 2015), a construct of TMEM16F, where the SCRD was replaced by the equivalent region of TMEM16A (TMEM16F^SCRD^), exhibited strongly compromised scrambling activity in our *in vitro* experiments (Figures 4D, 6F and S7I). On the contrary, the mutation Q623F, which removes a polar residue located on α6 facing the SCRD, did not affect lipid transport (Figures 4D and S7K). When investigating the TMEM16F^SCRD^ construct by electrophysiology, we observed robust currents with similar properties as WT and an increased potency for Ca^2+^ (Figures 6D and S7L). Thus, besides the point mutant R478A, also the SCRD chimera exerts different effects on lipid and ion permeation, indicating that both processes might be less coupled than previously anticipated (Jiang et al., 2017; Whitlock and Hartzell, 2016b; Yu et al., 2015).

## DISCUSSION

By combining data from cryo-electron microscopy, *in vitro* scrambling assays and electrophysiology, our study has revealed the basis for the diverse functional behavior of TMEM16F, a member of the TMEM16 family that facilitates the transport of lipids and ions across cellular membranes. Whereas the significance of lipid movement in a cellular environment is established, the role of ion conduction is currently unclear. Notably, due to the strong outward rectification, ionic currents would be small at resting potential but they could suffice to provide a shunt for the dissipation of charge during the scrambling of anionic lipids, which probably occurs at lower rates. Although our results offer detailed insights into potential transport mechanisms, the correspondence of the Ca^2+^-bound TMEM16F conformation to a defined functional state remains to some extend ambiguous. We thus envision potential alternative scenarios for ion and lipid permeation. In all scenarios, the structure of the Ca^2+^-free conformation describes an inactive conformation of the protein.

In the first scenario (‘out-of-the-groove mechanism’), the here reported calcium-bound TMEM16F structures represent a conformation close to the endpoint of the protein during activation (Figure 7A). Since, in the Ca^2+^-bound state the extracellular part of the subunit cavity remains shielded from the membrane, the diffusion of lipids would proceed outside of the cavity for about two-thirds of their transition. In this case, scrambling would be facilitated by interactions with the SCRD, which exposes polar residues to the inner leaflet of the bilayer at the gap between α-helices 4 and 6. Protein interactions with the lipid headgroups destabilize the membrane and consequently lower the barrier for lipid movement, which does not proceed in tight contact with the protein, as proposed in a recent study (Malvezzi et al., 2018). By contrast, ion conduction takes place in a sufficiently expanded protein-enclosed pore separated from the scrambling path. Upon activation, the observed movement of α-helix 6 towards α-helix 4 would change the structure in vicinity of the SCRD sufficiently to allow both, lipid movement and conduction. This scenario would imply that there are two classes of TMEM16 scramblases acting by distinct mechanisms. The first class would allow the passage of lipids through a membrane-accessible furrow and include fungal scramblases and their mammalian orthologues residing in intracellular organelles, such as TMEM16K. A second class, defined by TMEM16F and its paralogues, which are located at the plasma membrane, are closely related to the ion channels TMEM16A and B and thus show pronounced ion conduction properties but do not contain a membrane-accessible polar subunit cavity.

**Figure 7.**
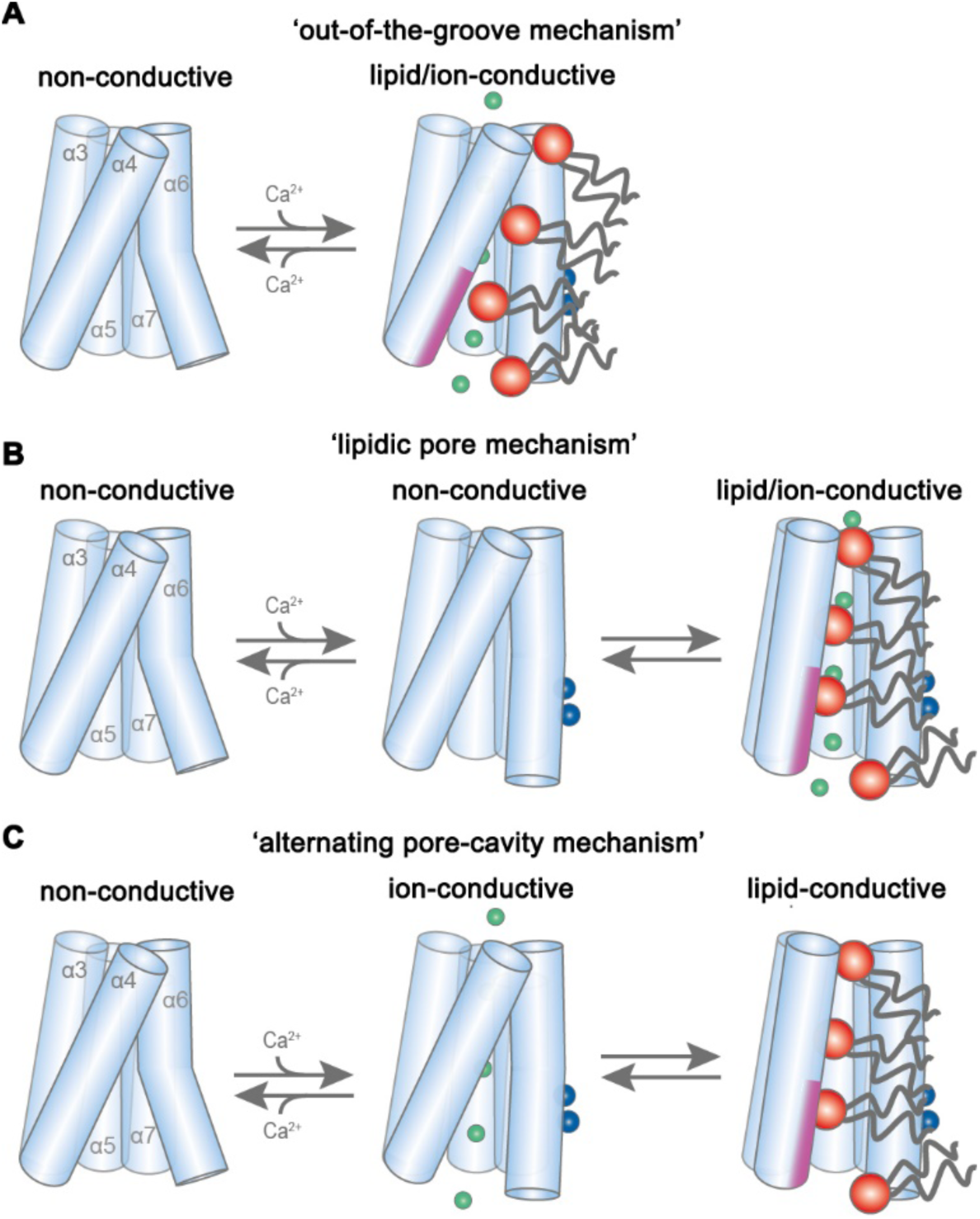
Potential transport mechanisms in TMEM16F. (A) ‘Out-of-the-groove mechanism’: upon calcium-activation the subunit cavity remains closed, ions and lipids permeate separate parts of the protein. (B) ‘Lipidic pore mechanism’: upon Ca^2+^ binding, the cavity opens with ions and lipids permeating the same open cavity conformation. (C) ‘Alternating pore-cavity mechanism’: After Ca^2+^-binding the open lipid-conductive cavity and ion conductive protein-surrounded pore are in equilibrium. This mechanism appears most likely in light of the functional data on TMEM16F and the results on a related study on the activation of the lipid scramblase nhTMEM16 (accompanying manuscript). (A to C) Schematic representation of TMEM16F α3–7 depicted as cylinders, bound calcium ions as blue spheres, conducting chloride ions as green spheres, scrambling lipid head groups as red spheres and their respective acyl chains as grey lines. The scrambling domain (SCRD) is indicated as purple area at the intracellular side of α-helix 4.

In a second scenario, scrambling would proceed on the surface of a subunit cavity that has opened to the membrane (Figures 7B and 7C). In this case, the ligand-bound TMEM16F structures display intermediates, whereby the conformational heterogeneity of α-helices 3 and 4 observed in the nanodisc samples indicates a potential rearrangement towards a fully active protein conformation. Consequently, the comparison to ligand-free structures reveals the first, ligand-dependent activation step. Full activation of the protein to a scrambling-competent state requires a consecutive transition that might be facilitated by lipid interaction with the protein presumably at the SCRD, which promotes a conformational change of α-helix 4 (Lee et al., 2018). This transition would lead to the exposure of a membrane-spanning hydrophilic furrow as observed in the scramblase nhTMEM16 (Brunner et al., 2014). This scenario is generally consistent with the activation mechanism for nhTMEM16, which is described in an accompanying manuscript. This fully active conformation was not observed in our structures, since it is either transient, as proposed for lipid scrambling in rhodopsin (Morra et al., 2018), or disfavored in our lipid nanodiscs, potentially due to a destabilizing lipid composition or the absence of interacting components (Schreiber et al., 2018; Ye et al., 2018). In such a scenario, we foresee two potential alternatives for the observed ion conduction. Conduction could either proceed in the same conformation as scrambling, in a pore that is partially lined by translocating lipids and thus be directly coupled to lipid movement (‘lipidic pore mechanism’), as suggested in previous investigations (Jiang et al., 2017; Whitlock and Hartzell, 2016b) (Figure 7B). Alternatively, ion conduction and scrambling could be mediated by distinct alternating conformations, which are at equilibrium in the Ca^2+^-bound state. Here, scrambling proceeds in an open subunit cavity, as observed for nhTMEM16 (accompanying manuscript), while ion conduction is catalyzed by a protein-enclosed pore that resembles the Ca^2+^-bound TMEM16F structures (‘alternating pore-cavity mechanism’, Figure 7C).

Whereas our structures do not show strong indication of an opened subunit cavity, the functional data are most consistent with distinct conformations facilitating ion conduction and scrambling as described in the ‘alternating pore-cavity mechanism’. In this case, mutations could shift the equilibrium between both states and have opposite effects on either function. This is observed for the mutants R478A and TMEM16F^SCRD^, which both mediate Ca^2+^-activated currents with increased ligand potency but show impaired lipid permeation properties. Additionally, this scenario accounts for the strong effect of the mutation of Arg 478 on lipid permeation (Figure 6E) (Gyobu et al., 2017), which in our structures is buried in the protein but would become exposed to the membrane in an open cavity. In contrast, the mutation of polar residues on the outside of the subunit cavity, which might contribute to scrambling in the ‘out-of-the-groove mechanism’, have little impact on lipid movement (Figures S7J and S7K). A shift in the equilibrium of the two states might also underlie the decrease in the open probability of the ion conduction pore at negative voltages. A definitive resolution of the mechanistic ambiguity will require further investigations, for which our study has provided an important foundation.

## SUPPLEMENTAL INFORMATION

Supplemental Information includes seven figures, one table and two movies and can be found with this article online at:

## ACKNOWLEDGMENTS

We thank S. Klauser, S. Rast and M. Punter for their help in establishing and maintaining the computer infrastructure, Y. Neldner for help during the generation of stable TMEM16F cell-lines and D. Deneka for advice with protein reconstitution into lipid nanodiscs. All members of the Dutzler and Paulino labs are acknowledged for their help at various stages of the project. This work was supported by a grant from the European Research Council (no 339116, AnoBest) to R.D. and by a grant from the NWO Start-Up (no 740.018.016) to C.P..

## AUTHOR CONTRIBUTIONS

C.A. generated stable cell-lines and expression constructs. C.A. and N.K.L. purified proteins for cryo-EM and functional characterization. C.A. reconstituted protein into nanodiscs and liposomes and carried out lipid transport experiments. N.K.L. recorded and analyzed electrophysiology data. C.P. prepared the samples for cryo-EM. V.C.M. G.T.O. and C.P. collected cryo-EM data. V.C.M and CP carried out image processing and C.P performed model building and refinement. C.A., N.K.L., V.C.M, R.D. and C.P. jointly planned experiments, analyzed the data and wrote the manuscript.

## METHODS

### Contact for Reagent and Resource Sharing

Further information and requests for resources and reagents should be directed to and will be fulfilled by the Lead Contact, Cristina Paulino.

### Cell lines

Flp-In™ T-REx™ 293 cell lines stably expressing TMEM16F were adapted to suspension cultures and were grown at 37 °C and 5% CO_2_ in EX-CELL 293 Serum-Free medium supplemented with 1% fetal bovine serum, 6 mM L-glutamine and 100 U/ml penicillin–streptomycin. GnTI^-^ cells were grown in HyClone HyCell TransFx-H medium supplemented with 1% fetal bovine serum, 4 mM L-glutamine, 1 g/l poloxamer 188 and 100 U/ml penicillin–streptomycin. Adherent HEK293T cells were grown in DMEM medium supplemented with 10% fetal bovine serum, 2 mM L-glutamine, 1 mM Sodium pyruvate and 100 U/ml penicillin–streptomycin.

### Construct preparation

The gene encoding murine TMEM16F (GenBank: BC060732) was obtained from Dharmacon-Horizon Discovery and stably inserted into a tetracycline-inducible HEK293T cell line using the Flp-In T-Rex System. For that purpose, the TMEM16F sequence was cloned into two pcDNA5/FRT expression vectors, one with C-terminal Rhinovirus 3C protease recognition site, Myc and Streptavidin-binding peptide (SBP) tags and the other with an extra Venus tag before the Myc and SBP tags. Both cell lines were created in the same manner: Briefly, 1 µg of TMEM16F expression vector and 9 µg pOG44 recombinase were co-transfected after mixing with 30 µg Fugene 6 transfection reagent. Hygromycin (50 µg/ml in the first week and 100 µg/ml afterwards) was used as selection marker and resistant foci appeared after 2 weeks. Per cell line, 15 foci were expanded and TMEM16F expression was assayed after tetracycline induction (2 µg/ml) for 48 h, extraction with 2% *n* **-**dodecyl**-**β**-**d**-**maltopyranoside (DDM) and Western blot analysis using a primary anti-Myc antibody together with a secondary peroxidase-coupled antibody (Figure S1B).

For mutagenesis, the sequence of murine TMEM16F was cloned into FX-cloning compatible pcDNA3.1 vectors (Geertsma and Dutzler, 2011) and point mutations were introduced with the non-overlapping primers modified QuikChange method (Zheng et al., 2004). The sequence was not codon optimized except for the removal of a *SapI* restriction site. For experiments requiring protein purification, the sequence was cloned into an expression vector with C-terminal 3C recognition site, Myc and SBP tags. For electrophysiology experiments, an expression vector with Venus tag between the 3C site and Myc-SBP was used instead. All constructs were verified by sequencing. The TMEM16F^SCRD^ construct, with residues 525–559 mutated to their equivalent in murine TMEM16A, was synthesized by GenScript. The plasmid used for the expression of the membrane scaffold protein (MSP) 2N2 was obtained from Addgene (plasmid #29520).

### Protein expression and purification

Wild-type TMEM16F was expressed by tetracycline induction (2 µg/ml) of the stably transfected TMEM16F-3C-Myc-SBP cell line for 48–70h. After induction, the medium was further supplemented with 3.5 mM valproic acid. Mutant TMEM16F constructs were expressed by transient transfection of GnTI^−^ cells. For cell transfection, DNA was complexed in a 1:2.5 ratio with Polyethylenimine MAX 40 K in non-supplemented DMEM medium for 20 min before addition to the cells. After transfection, the medium was further supplemented with 3.5 mM valproic acid and the cells were collected after 48–60 h, washed with PBS and stored at -80 °C until further use.

Protein purification of wild-type and mutant TMEM16F was carried out at 4 °C and was completed within 14 h. In all cases, we have purified the protein under Ca^2+^-free conditions and added 1 mM Ca^2+^ when indicated during cryo-EM sample preparation. Cells were resuspended in 2% digitonin (PanReac AppliChem), 150 mM NaCl, 20 mM HEPES, pH 7.5, 5 mM EGTA and protease inhibitors and the membranes were solubilized by gentle mixing for 2 h. Solubilized proteins were isolated by centrifugation at 85 000 *g* for 30 min and allowed to bind to streptavidin UltraLink resin beads for 2 h under gentle agitation. The beads were loaded onto a gravity column and washed with 60 column volumes (CV) of SEC buffer containing 0.1% digitonin (EMD Millipore), 150 mM NaCl, 20 mM HEPES, pH 7.5 and 2 mM EGTA. Bound protein was eluted with 3 CV of SEC buffer supplemented with 4 mM biotin. Samples used for Cryo-EM were digested with PNGaseF for 2 h. Protein was concentrated using a 100 kDa cutoff filter, filtered through a 0.22 µm filter and loaded onto a Superose 6 10/300 GL column pre-equilibrated with SEC buffer. Protein-containing peak fractions were pooled, concentrated, filtered and immediately used for either Cryo-EM sample preparation or reconstitution into nanodiscs or liposomes. The MSP 2N2 protein was expressed and purified as described (Ritchie et al., 2009). In this case, the N-terminal polyhistidine tag was not cleaved after purification.

Nanodisc reconstitution was performed as described (Gao et al., 2016) with minor modifications. Purified protein was incorporated into nanodiscs composed of a1-palmitoyl-2-oleoyl-glycero-3-phosphocholine (POPC) and 1-palmitoyl-2-oleoyl-sn-glycero-3-phospho-(1'-rac-glycerol) (POPG) mixture at a molar ratio of 3:1. After mixing, the chloroform-dissolved lipids were initially dried under a nitrogen stream, washed with diethyl ether and again dried under nitrogen stream and vacuum desiccation overnight. The lipid mix was solubilized in 10% DDM, at a final lipid concentration of 10 mM. A molar ratio of 2:10:2200 of TMEM16F:MSP 2N2:lipids was used for reconstitution. Purified protein was incubated with the lipid mix on ice for 40 min. Subsequently, 2N2 was added to the mixture and further incubated for 40 min on ice. Detergent was removed overnight after addition of 50 mg of SM-II biobeads per mg of DDM. Biobeads were removed from the clear sample by column filtration and the nanodisc suspension was incubated with streptavidin UltraLink resin for 2 h. Further purification proceeded as described for the digitonin-solubilized protein, except that no detergent was present in any of the buffers.

### Liposome reconstitution and scrambling experiments

For the characterization of lipid scrambling, purified TMEM16F was reconstituted into liposomes containing trace amounts (0.5% weight/weight) of fluorescently (nitrobenzoxadiazole, NBD) labeled lipids. The lipid mix used for the majority of scrambling experiments was soybean polar lipids extract with 20% cholesterol (mol/mol) and 0.5% 18:1-06:0 NBD-PE. To investigate whether scrambling proceeds independently of the chemical nature of the headgroup of the labeled lipid or the location of the fluorophore, the tail labeled 18:1-06:0 NBD-PE was substituted by 18:1-06:0 NBD-PS or head-labeled 14:0 NBD-PE while maintaining the other liposome components unchanged. To characterize the protein activity in the lipid mix used for the preparation of nanodisc samples, we reconstituted TMEM16F into liposomes composed of 3 POPC : 1 POPG : 0.5 % 18:1-06:0 NBD-PE. Lipids were treated similarly as the ones used for nanodisc reconstitution until solubilization. Liposomes were prepared as described (Geertsma et al., 2008). For liposomes, the lipids were solubilized in 20 mM HEPES pH 7.5, 300 mM KCl and 2 mM EGTA (buffer A) in a final concentration of 20 mg/ml, sonicated, subjected to three freeze-thaw cycles in liquid N2 and stored at -80 °C. The liposomes were protected from direct light as much as possible. Prior to reconstitution, the lipids were extruded 21 times through a 400 nm pore polycarbonate membrane and diluted to 4 mg/ml in buffer A. Liposomes were destabilized by addition of Triton X-100 aliquots in 0.02% steps until the onset of lipid solubilization monitored by the decrease of the scattering at 540 nm in a spectrophotometer. After addition of another aliquot of 0.1% Triton X-100, the protein was added at a lipid-to-protein ratio of 100:1 (weight/weight) unless stated otherwise to the destabilized liposomes and the mixture was incubated at room temperature (RT) for 15 min under gentle agitation. Subsequently, 20 mg of SM-II biobeads per mg of lipids were added four times to completely remove digitonin and Triton X-100. After 45 min, the sample was moved to 4 °C. Proteoliposomes were harvested after 24 hours by filtration to remove the biobeads followed by centrifugation at 150,000 *g* for 30 min. The pelleted proteoliposomes were resuspended in buffer A with solutions containing the appropriate amount of Ca(NO3)2 to obtain the indicated concentrations of free Ca^2+^ calculated with the onlineWEBMAXC calculator (Bers et al., 2010). The liposomes were resuspended at either 10 or 20 mg/ml and subjected to three freeze-thaw cycles in liquid N2. All remaining steps were carried out at RT. After extrusion through a 400 nm membrane pre-equilibrated in buffer A containing the desired amount of free Ca^2+^, proteoliposomes were diluted to 0.2 mg/ml in 80 mM HEPES pH 7.5, 300 mM KCl, 2 mM EGTA plus Ca^2+^ (buffer B). Scrambling was performed essentially as described for other scramblases of the TMEM16 family (Brunner et al., 2014; Malvezzi et al., 2013). Scrambling was monitored by spectrofluorimeter measurements of the NBD fluorophore with an excitation wavelength of 470 nm and emission wavelength of 530 nm. After 60 s of recording, 30 mM of the membrane-impermeable reducing agent sodium dithionite was added to irreversibly bleach exposed NBD groups. The total recording time was of 400 seconds unless stated otherwise and the sample was stirred during the entire measurement. The NBD-fluorescence decay was plotted as F/Fmax. To investigate the ligand-dependence of the scrambling rate, we measured the NBD-fluorescence value 120 s after the addition of dithionite at different Ca^2+^ concentrations and quantified scrambling activity as 1-F_Ca2_+/F0_Ca2_+ since we were unable to detect any pronounced scrambling activity for TMEM16F in the absence of Ca^2+^. The effect of Ca^2+^ on the scrambling rate was studied using symmetric buffer conditions, with the same Ca^2+^ concentration inside and outside the liposomes. To investigate the delay time between calcium addition and scrambling activation we reconstituted TMEM16F in calcium-free liposomes and recorded the NBD-fluorescence decay upon addition of 100 µM Ca(NO_3_)_2_ to the outside buffer, 120 s after dithionite addition, and monitored fluorescence for extra 420 s. The same batch of liposomes was incubated in buffer B containing 100 µM Ca(NO_3_)_2_ for 10 minutes prior to measurement as control.

In empty liposomes, addition of dithionite causes a decrease of the fluorescence to about half of its initial value, due to the bleaching of the fraction of fluorescent lipids located in the outer leaflet of the bilayer (Figure S1D). In contrast, proteoliposomes containing an active scramblase show a stronger time-dependent decrease of the fluorescence to a plateau, which corresponds to the fraction of fluorescent lipids that are not accessible to dithionite since they either reside in liposomes not containing a reconstituted scramblase or they are located on the inside of a multi-lamellar vesicle (Malvezzi et al., 2018; Ploier and Menon, 2016) (Figures 1A, S1D-1G). For different reconstitutions, we find similar plateau values for empty liposomes and proteoliposomes in Ca^2+^-free conditions. In contrast, the plateau of fully activated samples is consistent for different constructs using the same batch of destabilized lipids but it varies between different reconstitutions, with fluorescence levels ranging from 18% to 30%. We attribute this difference to the efficiency of reconstitution that is dependent on the batch of destabilized liposomes. Independent of the plateau, all reconstitutions showed a very similar Ca^2+^-dependence. Due to the variable reconstitution efficiency, all the mutants were purified in parallel with wild type (WT) TMEM16F and reconstituted using the same batch of lipids in equivalent reconstitution conditions. Protein incorporation into liposomes for WT and mutants was verified by Western blot against the Myc tag after solubilization of proteoliposomes with DDM. With the exception of TMEM16F^SCRD^, all mutants reconstituted with an efficiency similar to WT (Figure S7I). TMEM16F^SCRD^ behaved well during purification but reconstituted with somewhat lower efficiency. For that purpose, it was compared to WT reconstituted at lower lipid to protein ratio of 150:1. For this case proteoliposomes contained similar protein levels (Figure S7I).

### Electrophysiology

For electrophysiology, HEK293T cells were transfected with plasmids at a concentration of 8 µg DNA per 100 mm culture dish with 25 µl FuGENE 6 transfection reagent. Transfected cells were identified by Venus fluorescence and used for patch clamp experiments within 24–72 h of transfection. All recordings were performed in the inside-out configuration (Hamill et al., 1981) at RT (20–22 °C). Inside-out patches were excised from HEK293T cells expressing the TMEM16F construct of interest after the formation of a gigaohm seal. Seal resistance was typically 4–8 GΩ or higher. Patch pipettes were pulled from borosilicate glass capillaries (OD 1.5 mm, ID 0.86 mm) and were fire-polished using a microforge. Pipette resistance was typically 3–8 MΩ when filled with pipette solutions. Voltage-clamp recordings were performed using the Axopatch 200B amplifier controlled by the Clampex 10.6 software through Digidata 1440. The data were sampled at 10 kHz and filtered at 1 kHz. Liquid junction potential was not corrected. Step-like solution exchange was elicited by analogue voltage signals delivered through Digidata 1440. All recordings were measured in symmetrical NaCl solutions except for selectivity experiments. The background current was recorded in Ca^2+^ free solution and substracted prior to analysis. Ca^2+^-free intracellular solution contained 150 mM NaCl, 5 mM EGTA, and 10 mM HEPES, pH 7.40. High intracellular Ca^2+^ solution, with a free concentration of 1 mM, contained 150 mM NaCl, 5.99 mM Ca(OH)_2_, 5 mM EGTA, and 10 mM HEPES, pH 7.40. The pH was adjusted using 1 M NMDG-OH solution. Intermediate Ca^2+^ solutions were obtained by mixing high Ca^2+^ and Ca^2+^-free intracellular solutions at the ratio calculated according to the WEBMAXC calculator (Bers et al., 2010) free Ca^2+^ concentrations above 1 mM (up to 10 mM) was adjusted by adding CaCl_2_ from a 1 M stock solution. The Ca^2+^-free intracellular solution was also used as the pipette solution. For permeability experiments using (NMDG)_2_SO_4_ to compensate for the ionic strength, the appropriate NaCl concentration were adjusted by mixing NaCl stock solutions and (NMDG)_2_SO_4_ stock solutions at the required ratios. Stock NaCl solutions were the same as above. Stock (NMDG)_2_SO_4_ solutions contained 100 mM (NMDG)_2_SO_4_, 5.99 mM Ca(OH)_2_, 5 mM EGTA, 10 mM HEPES, pH 7.40, and 100 mM (NMDG)_2_SO_4_, 5 mM EGTA, and 10 mM HEPES, pH 7.40.

Current of TMEM16F runs down in the excised patch configuration. In order to obtain a more accurate EC_50_ of Ca^2+^-activation, rundown correction was performed using a previously described method (Lim et al., 2016). In short, a reference Ca^2+^ pulse was applied at a regular time interval before and after the test pulse. The magnitude of the test pulse was normalized to the average magnitude of the pre- and post-reference pulses. The normalized concentration-response data were fitted to the Hill equation. To correct for rundown during experiments used for the quantification of rectification, the instantaneous and steady-state current at each voltage step was divided by the ratio of the remaining current at the 80 mV pre-pulse and expressed as normalized current (I/I80mV). Statistical difference between the constructs were determined with one-way ANOVA and Tukey-Kramer post-hoc test. Values were considered significantly different if *P* < 0.05. By this criterion, all the shifts in the EC_50_ for mutants described in the study except for the mutant R478A were determined as significant. The shift in R478A is slightly above the significance threshold. For measurements of rectification, only the shift of the steady state currents of the mutant R478A was significant.

### Cryo-electron microscopy sample preparation and imaging

2.5 μl of freshly purified protein at a concentration of 3.3 mg ml^−1^ when solubilized in digitonin and 1 mg ml^−1^ when reconstituted in nanodiscs, were applied on holey-carbon cryo-EM grids (Quantifoil Au R1.2/1.3, 200, 300 and 400 mesh), which were prior glow-discharged at 5 mA for 30 s. For the datasets obtained for the Ca^2+^-bound structures, samples were supplemented with 1 mM CaCl2 before freezing. Grids were blotted for 2-5 s in a Vitrobot (Mark IV, Thermo Fisher) at 10-15 °C temperature and 100% humidity, subsequently plunge-frozen in liquid ethane and stored in liquid nitrogen until further use. Cryo-EM data were collected on a 200 keV Talos Arctica microscope (Thermo Fisher) using a post-column energy filter (Gatan) in zero-loss mode, a 20 eV slit, a 100 µm objective aperture, in an automated fashion using EPU software (Thermo Fisher) on a K2 summit detector (Gatan) in counting mode. Cryo-EM images were acquired at a pixel size of 1.012 Å (calibrated magnification of 49,407x), a defocus range from –0.5 to –2 µm, an exposure time of 9 s with a sub-frame exposure time of 150 ms (60 frames), and a total electron exposure on the specimen of about 52 electrons per Å^2^. Best regions on the grid were manually screened and selected with the help of an in-house script to calculate the ice thickness. Data quality was monitored on the fly using the software FOCUS (Biyani et al., 2017).

### Image Processing

For the detergent dataset collected in presence of Ca^2+^, the 5633 dose-fractionated cryo-EM images recorded (final pixel size 1.012 Å) were subjected to motion-correction and dose-weighting of frames by MotionCor2 (Zheng et al., 2017). The CTF parameters were estimated on the movie frames by ctffind4.1 (Rohou and Grigorieff, 2015). Bad images showing contamination, a defocus above –0.5 or below –2 µm, or a bad CTF estimation, were discarded. The resulting 4225 images were used for further analysis with the software package RELION2.1 (Kimanius et al., 2016). Particles were initially picked automatically from a subset of the dataset using 2D-class averages from the previously obtained TMEM16A cryo-EM map (Paulino et al., 2017a) as reference. The resulting particles where used to create a better reference for autopicking, which was then repeated on the whole dataset. The final round of autopicking yielded 1,348,247 particles, which were extracted with a box size of 220 pixels, and initial classification steps were performed with two-fold binned data. False positives were removed in the first round of 2D classification. Remaining particles were subjected to several rounds of 2D classification, resulting in 680,465 particles that were further sorted in several rounds of 3D classification. The TMEM16A cryo-EM map, low-pass filtered to 50 Å, was used as a reference for the first round of 3D classification and the best output class was used in subsequent jobs in an iterative way. The best 3D classes, comprising 219,302 particles, were subjected to auto-refinement, yielding a map with a resolution of 3.8 Å. In the last refinement iteration, a mask excluding the micelle was used and the refinement was continued until convergence, which improved the resolution up to 3.5 Å. Using a mask generated from the final PDB model, we obtained a map at 3.4 Å. Finally, the newly available algorithms for CTF refinement and Bayesian polishing implemented in Relion3.0 (Zivanov et al., 2018), where applied to further improve the resolution to 3.18 Å with a map sharpened at –94 Å^2^. During final 3D classification and auto-refinement jobs, a C2-symmetry was imposed. Local resolution was estimated by RELION. All resolutions were estimated using the 0.143 cut-off criterion (Rosenthal and Henderson, 2003) with gold-standard Fourier shell correlation (FSC) between two independently refined half maps (Scheres and Chen, 2012). During post-processing, the approach of high-resolution noise substitution was used to correct for convolution effects of real-space masking on the FSC curve (Chen et al., 2013). The directional resolution anisotropy of density maps was quantitatively evaluated using 3DFSC (Tan et al., 2017).

For the other datasets, a similar workflow for image processing was applied. In case of the detergent dataset collected in absence of Ca^2+^, a total of 1,314,676 particles were extracted after auto-picking from 4,621 images. Several rounds of 2D and 3D classification resulted in a final number of 194,284 particles, which, after refinement yielded a 4.1 Å map. Providing a mask in the last iteration of the refinement, the resolution was improved to 3.8 Å. After post-processing the resolution improved to 3.6 Å. For the dataset in nanodiscs in the presence of Ca^2+^, 1,019,012 particles were picked from 4,480 images and extracted with a box size of 256 pixels. The final set of 186,487 particles was refined, yielding a map with a resolution of 3.9 Å, which after postprocessing was improved to 3.5 Å. Finally, the dataset in nanodisc in absence of Ca^2+^ resulted in 1,593,115 auto-picked particles from 6,465 images, which were reduced to 280,891 particles after several rounds of 2D and 3D classification. Refinement was performed providing a mask in the last iteration, reaching a resolution of 3.3 Å after post-processing.

The analysis of structural heterogeneity within each dataset was conducted by running an additional 3D classification step with finer angular sampling on the final particle set. While some structural flexibility was revealed for α3 and α4 in the dataset in presence of Ca^2+^ (Figures S2I and S4I), no alternative conformations could be identified in the dataset in absence of Ca^2+^ (Figures S3I and S5I), as reflected in the better resolved density of the helices (Figure S6A).

Finally, as revealed by the angular distribution and the 3D FSC plots obtained for the data in nanodiscs, the samples suffered from a favorite orientation of the particles in vitreous ice, thereby compromising to some extent the detailed features of the maps (Figures S4, S5 and S6A). The problem is less pronounced in the nanodisc data obtained in presence of Ca^2+^ than in the dataset of the Ca^2+^-free sample. Nonetheless, the data in nanodiscs are of sufficient quality to allow for a comparison of corresponding states of TMEM16F in detergent and a lipid environment.

### Model building refinement and validation

The model of the Ca^2+^-bound state of mTMEM16F was built in COOT (Emsley and Cowtan, 2004), using the structure of TMEM16A (PDB 5YOB) as template. The density was of sufficiently high resolution to unambiguously place the model consisting of residues 198-222, 240-423, 448-486, 503-637, 647-788 and 798-875. Additional density at the N-terminus encompassing residues 116-125 and 134-192 could be interpreted as poly-alanine chain with ambiguous register. The model was improved by real-space refinement in Phenix (Adams et al., 2010), whereby secondary structure elements and the symmetry between both subunits of the dimeric protein were constrained. Coordinates were manually edited in COOT after each refinement cycle. The structure of the Ca^2+^-free structure in digitonin and both Ca^2+^-free and Ca^2+^-bound structure obtained in nanodiscs of mTMEM16F were built using the refined Ca^2+^-bound structure as starting model. Major conformational changes were restricted to the second half of α-helix 6 and to some extent to α-helix 3 and 4. These regions were rebuilt manually in COOT and subsequently refined in Phenix, as described above. Due to the strong anisotropy of the dataset of TMEM16F obtained in nanodiscs in absence of calcium, only the transmembrane domain and some sections of the extracellular loops were built. For validation of the refinement, Fourier shell correlations (FSC) between the refined model and the final map were determined (FSC_SUM_, Figures S2E, S3E, S4E and S5E). To monitor the effects of potential over-fitting, random shifts (up to 0.3 Å) were introduced into the coordinates of the final model, followed by refinement in Phenix against the first unfiltered half-map. The FSC between this shaken-refined model and the first half-map used during validation refinement is termed FSC_work_, the FSC against the second half-map, which was not used at any point during refinement, FSC_free_. The marginal gap between the curves describing FSC_work_ and FSC_free_ indicates no over-fitting of the model. The higher resolution of α-helix 3 obtained in the cryo-EM map of TMEM16F in digitonin in absence of Ca^2+^ allowed the unambiguous assignment of its residues. Based on this structure and the conservation between both paralogs, we remodeled α3 of TMEM16A (Paulino et al., 2017a), which in its original data was poorly resolved. In the new model of TMEM16A, the register of α3 has shifted by three residues and Arg 515 now superimposes with Arg 478 in TMEM16F, both pointing into the pore lumen (Figure 4B). The corrected model of TMEM16A was used for structural comparisons throughout the manuscript. The pore diameter was calculated in HOLE (Smart et al., 1996). Pictures were generated using pymol (The PyMOL Molecular Graphics System, Version 2.0 Schrödinger, LLC), dino (http://www.dino3d.org), chimera (Pettersen et al., 2004) and chimeraX (Goddard et al., 2018).

### Electrostatic calculations

The electrostatic potential was calculated as described (Paulino et al., 2017a). The linearized Poisson-Boltzmann equation was solved in CHARMM (Brooks et al., 1983; Im et al., 1998) on a 200 Å × 140 Å × 190 Å grid (1 Å grid spacing) followed by focusing on a 135 Å x 100 Å x 125 Å grid (0.5 Å grid spacing). Partial protein charges were derived from the CHARMM36 all-hydrogen atom force field. Hydrogen positions were generated in CHARMM. The protein was assigned a dielectric constant (*ϵ*) of 2. Its transmembrane region was embedded in a 30-Å-thick slab (*ϵ* = 2) representing the hydrophobic core of the membrane and two adjacent 15-Å-thick regions (*ϵ*=30) representing the headgroups. This region contained a cylindrical hole around the water-filled intracellular vestibule of one subunit and was surrounded by an aqueous environment (*ϵ* =80) containing 150 mM of monovalent mobile ions.

### Statistics and Reproducibility

Electrophysiology data were repeated multiple times from different transfections with very similar results. Conclusions of experiments were not changed upon inclusion of further data. In all cases, leaky patches were discarded. Statistical difference between the constructs were determined with one-way ANOVA and Tukey-Kramer post-hoc test.

### Data availability

The three-dimensional cryo-EM density maps of calcium-bound mTMEM16F in detergent and nanodiscs have been deposited in the Electron Microscopy Data Bank under accession numbers EMD-XXX and EMD-XXX, respectively. The maps of calcium-free samples in detergent and nanodiscs were deposited under accession numbers EMD-XXX and EMD-XXX, respectively. The deposition includes the cryo-EM maps, both half-maps, and the mask used for final FSC calculation. Coordinates of all models have been deposited in the Protein Data Bank under accession numbers XXX (Ca^2+^-bound, detergent), XXX (Ca^2+^-bound, nanodisc), XXX (Ca^2+^-free, detergent) and XXX (Ca^2+^-free, nanodisc). Data can be made available upon request.

## SUPPLEMENTAL INFORMATION

**Figure S1.**
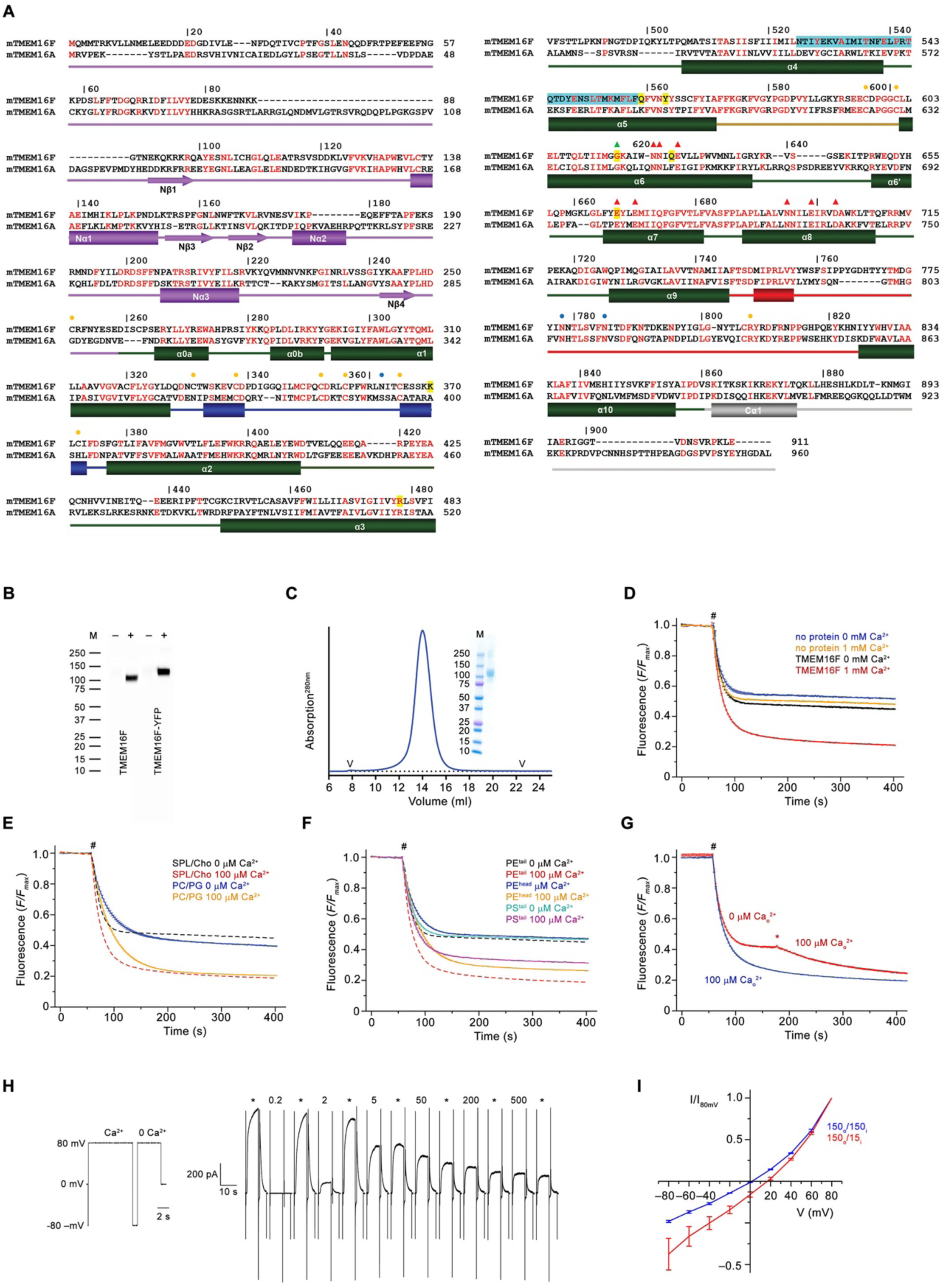
Sequence alignment, biochemistry and functional characterization, Related to Figure 1. (A) Protein sequences of murine TMEM16F (UniProt: Q6P9J9.1) and murine TMEM16A (UniProt Q8BHY3.2) were aligned with Clustal Omega (Sievers et al., 2011). Numbering corresponds to TMEM16F. Secondary structure elements: green, transmembrane domain; violet, N-terminal domain; grey, C-terminal region; blue, α1–α2 loop; beige, α5–α6 loop; red, α9–α10 loop). Boundary of secondary structure elements refers to TMEM16F, except for the N-terminal domain, which corresponds to TMEM16A due to uncertainty in the register of TMEM16F in this region. Orange circles mark the position of cysteines involved in disulfide bonds, blue circles mark putative sites of glycosylation indicated by residual density. Red triangles correspond to residues of the Ca^2+^-binding site and the green triangle to the hinge for conformational rearrangements in α6. The SCRD is highlighted in cyan and positions mutated in this study in yellow. (B) Western blot of stable cell-lines expressingTMEM16F before (−) and after (+) induction with tetracycline. TMEM16F and a c-terminal YFP fusion (TMEM16F-YFP) were detected with an anti-myc antibody. (C) Size-exclusion chromatogram of purified TMEM16F. V indicates void and dead volumes of the column. Inset shows SDS-Page of peak fractions. The molecular weight marker (M) and corresponding masses (kDa) are indicated. (D) Fluorescence-based lipid transport assay in proteoliposomes containing soybean polar lipids and cholesterol used throughout the study. The fluorescence of tail-labeled NBD-PE was quenched by addition of dithionite (#) to the outside, which leads to a reduction of the fluorescence to about half in liposomes not containing any protein. The reduction in TMEM16F-containing proteoliposomes in the absence of Ca^2+^ underlines the poor activity of the protein in absence of its ligand. The stronger decrease at high Ca^2+^ concentration reflects activity of TMEM16F as lipid scramblase. The fluorescence at the plateau is due to lipids that are not accessible to dithionite. (E) Comparison of TMEM16F scrambling activity in proteoliposomes of different lipid composition. Solid lines depict data from proteoliposomes containing a PC/PG mixture used for reconstitution into nanodiscs, which show a Ca^2+^-activated activity that proceeds with slower kinetics and larger basal activity compared to soybean and cholesterol containing liposomes (c, dashed line). (F) Scrambling of proteoliposomes with different labeled lipids. TMEM16F is capable of scrambling tail-labeled NBD-PS (PS^tail^) and head-labeled NBD-PE (PE^head^), thus underlining its poor substrate specificity. Data for tail-labeled NBD-PE used for most experiments (c, dashed line) are shown for comparison. (G) Instantaneous activation of TMEM16F is shown by addition of Ca^2+^ (*) 120 s after the addition of dithionite (#), which causes a further decrease of the fluorescence due to the exposure of labeled lipid that are transported from the inner leaflet following the activation of the scramblase (red). Liposomes that were pre-incubated with Ca^2+^ on their outside before addition of dithionite are shown in blue for comparison. (D–G) Traces show average of three experiments from the same reconstitution. Errors are s.e.m.. # indicates addition of dithionite. (H) Representative current response of TMEM16F at different Ca^2+^ concentrations measured in excised inside-out patches. Numbers indicate Ca^2+^ concentration (µM), * indicates reference pulses recorded at 20 µM Ca^2+^ used for rundown correction. Individual sweeps are separated by a regular time interval (15–18 s) and are concatenated to form a continuous trace. Inset (left) shows voltage protocol used for each sweep in all concentration-response relationships throughout the study. (I) Normalized I-V relationships of TMEM16F at symmetric (blue) and in a 10-fold gradient of NaCl concentration (red). The osmolarity was adjusted by addition of NMDG-sulfate. The shift in the reversal potential in asymmetric conditions indicates a slight selectivity for cations over anions. Data show averages of three biological replicates, errors are s.e.m..

**Figure S2.**
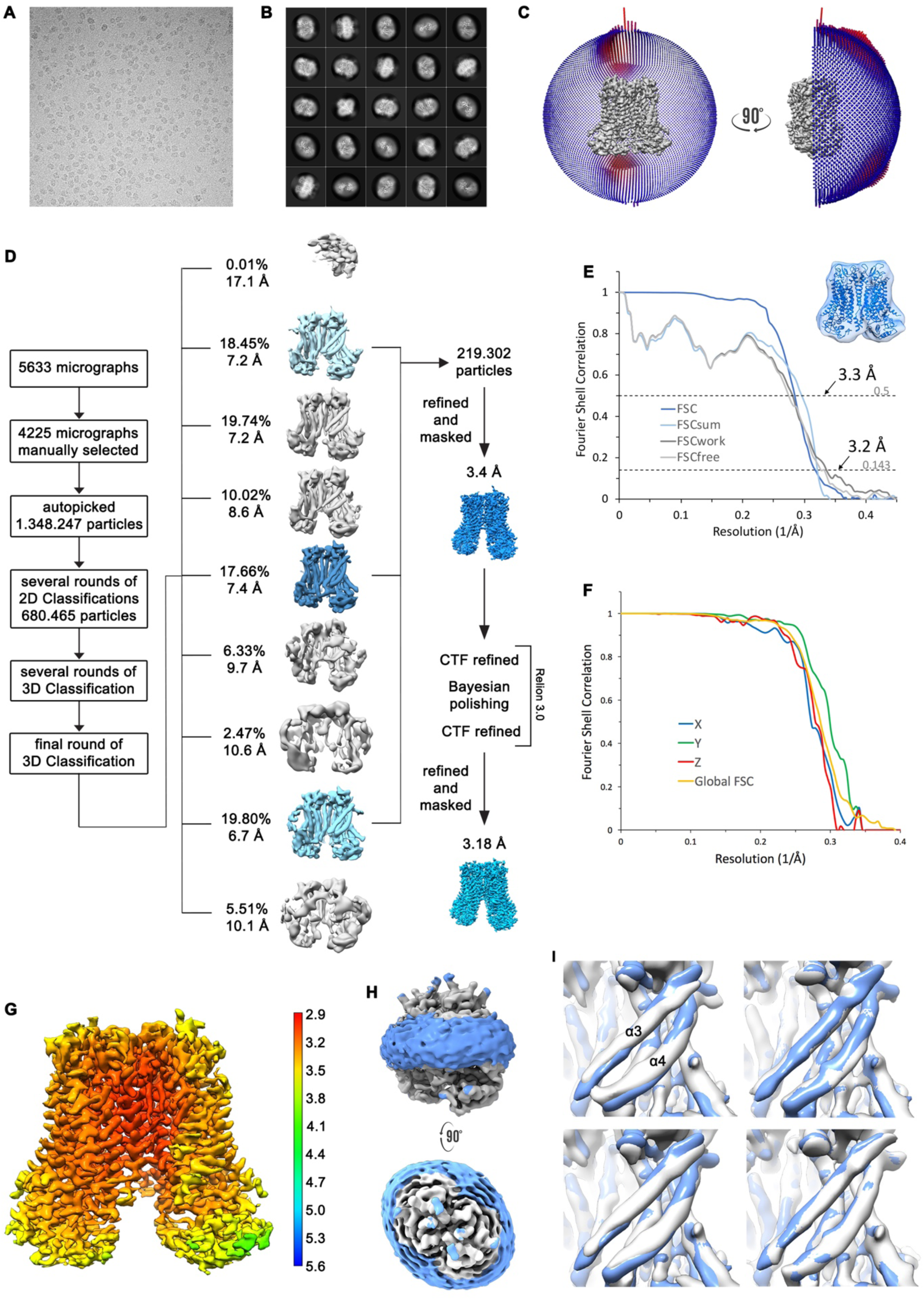
Structure Determination of TMEM16F in complex with Ca^2+^ in digitonin, Related to Figure 2. (A) Representative cryo-EM image and (B) 2D-class averages of vitrified mTMEM16F in a Ca^2+^-bound state in digitonin. (C) Angular distribution plot of particles included in the final C2-symmetric 3D reconstruction. The number of particles with their respective orientation is represented by length and color of the cylinders. (D) Image processing workflow. (E) FSC plot used for resolution estimation and model validation. The gold-standard FSC plot between two separately refined half-maps is shown in dark blue and indicates a final resolution of 3.18 Å. FSC validation curves for FSC_sum_, FSC_work_ and FSC_free_, as described in Methods, are shown in light blue, dark grey and light grey, respectively. A thumbnail of the mask used for FSC calculation overlaid on the atomic model is shown in the upper right corner and thresholds used for FSC_sum_ of 0.5 and for FSC, FSC_work_ and FSC_free_ of 0.143 are shown as dashed lines. (F) Anisotropy estimation plot of the final map. The global FSC curve is represented in yellow. The directional FSCs along the x, y and z axis displayed in blue, green and red, respectively, are indicative for an isotropic dataset. (G) Final reconstruction map colored by local resolution as estimated by Relion, indicate regions of higher resolution. (H) Representation of the map including the detergent micelle (low-pass filtered to 6.5 Å) reveals no noticeable distortion of the micelle (blue). (I) Analysis of structural heterogeneity by subjecting the final dataset to another round of 3D classification, reveals alternative conformations of α3 and α4. Shown are superpositions of obtained 3D classes (grey) on the final cryo-EM map (blue).

**Figure S3.**
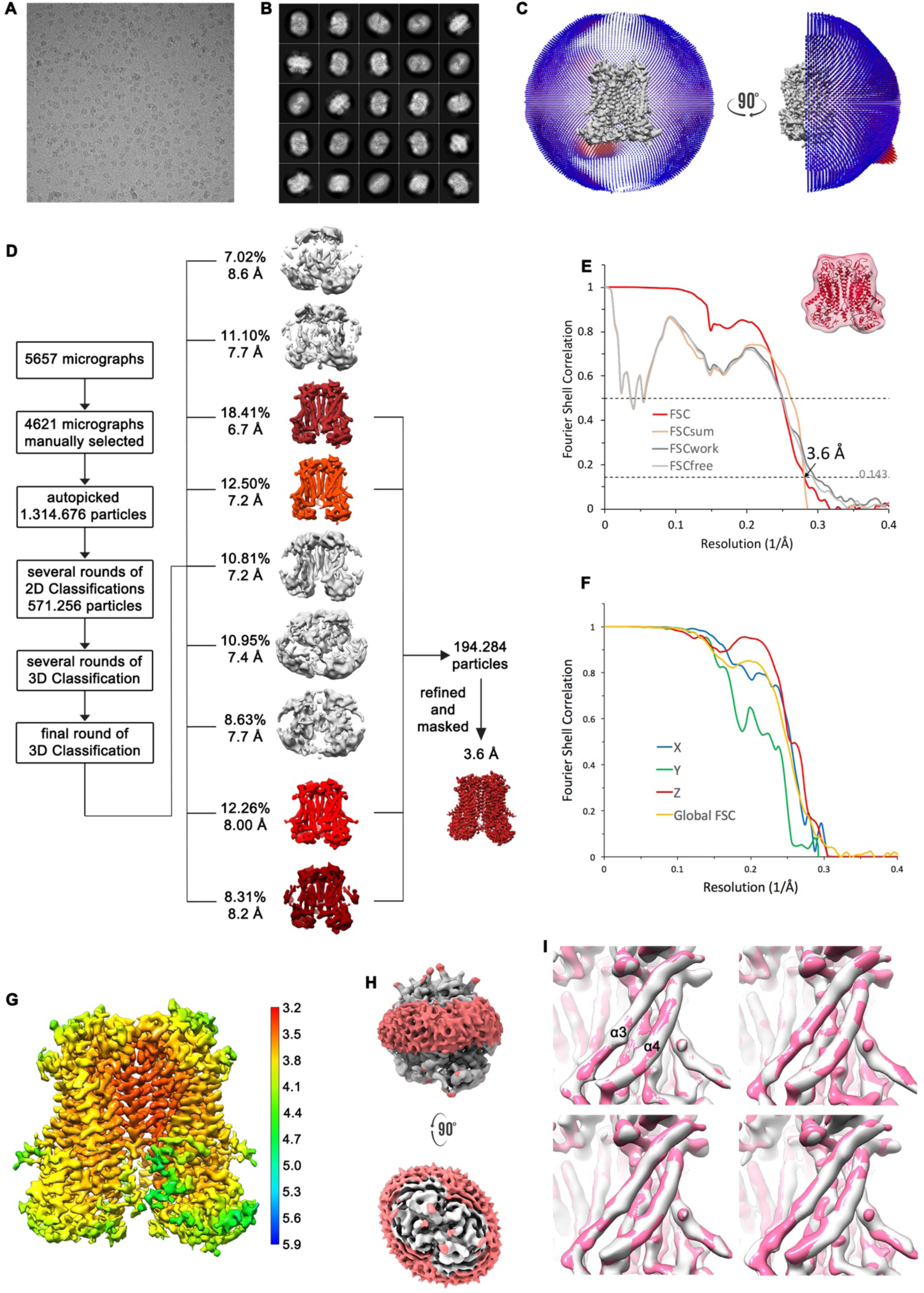
Structure Determination of TMEM16F in absence of Ca^2+^ in digitonin, Related to Figure 2. (A) Representative cryo-EM image and (B) 2D-class averages of vitrified mTMEM16F in a Ca^2+^-free state in digitonin. (C) Angular distribution plot of particles included in the final C2-symmetric 3D reconstruction. The number of particles with their respective orientation is represented by length and color of the cylinders. (D) Image processing workflow. (E) FSC plot used for resolution estimation and model validation. The gold-standard FSC plot between two separately refined half-maps is shown in red and indicates a final resolution of 3.6 Å. FSC validation curves for FSC_sum_, FSC_work_ and FSC_free_, as described in Methods, are shown in orange, dark grey and light grey, respectively. A thumbnail of the mask used for FSC calculation overlaid on the atomic model is shown in the upper right corner and threshold used for FSC_sum_ of 0.5 and for FSC, FSC_work_ and FSC_free_ of 0.143 are shown as dashed lines. (F) Anisotropy estimation plot of the final map. The global FSC curve is represented in yellow. The directional FSCs along the x, y and z axis shown in blue, green and red, respectively, are indicative for a largely isotropic dataset. (G) Final reconstruction map colored by local resolution as estimated by Relion, indicating regions of higher resolution. (H) Representation of the map including the detergent micelle (low-pass filtered to 6.5 Å) reveals no noticeable distortion of the micelle (red). (I) Analysis of structural heterogeneity by subjecting the final dataset to another round of 3D classification, reveals no alternative conformations of α3 and α4. Shown are superpositions of obtained 3D classes (grey) on the final cryo-EM map (magenta).

**Figure S4.**
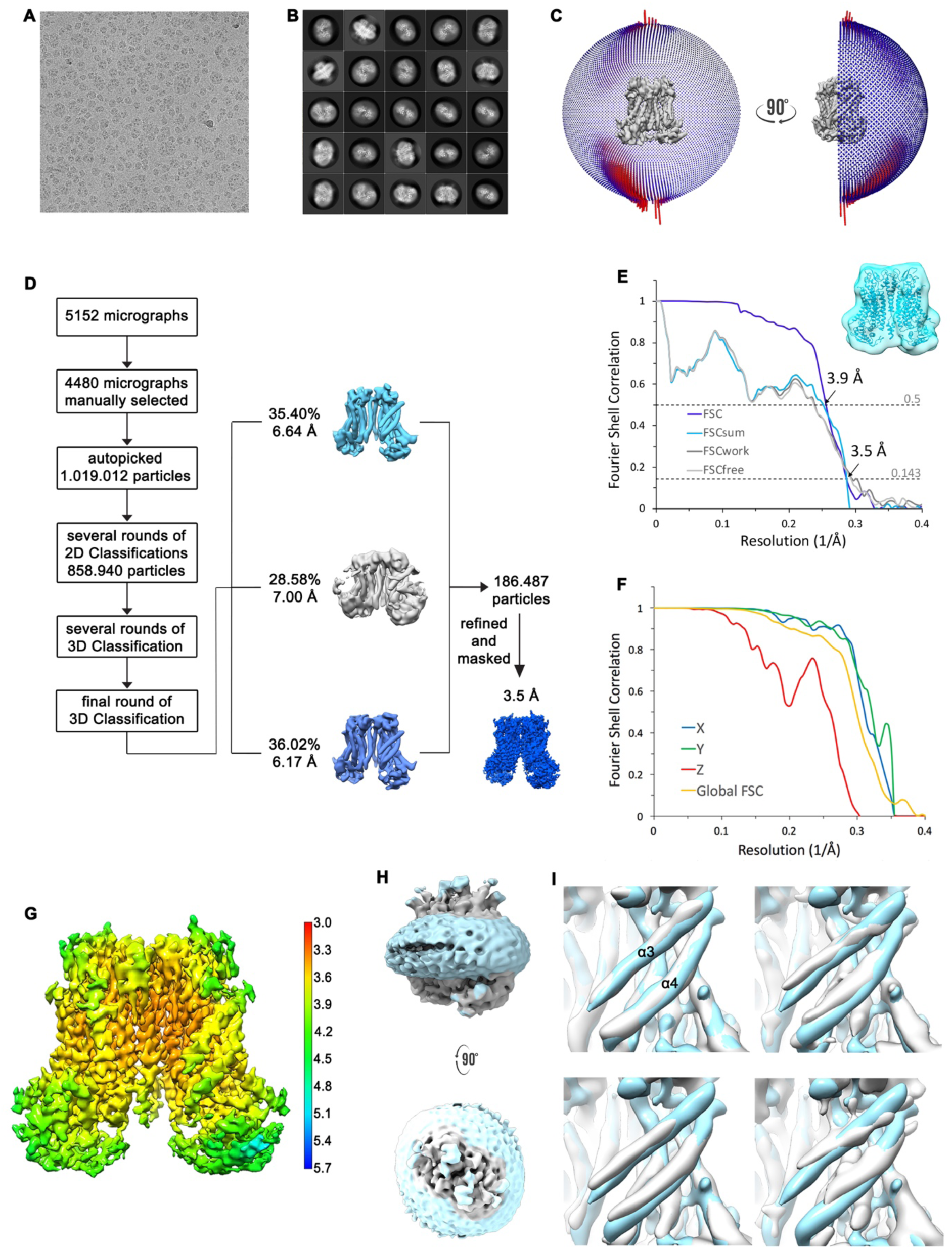
Structure Determination of TMEM16F in complex with Ca^2+^ in nanodisc, Related to Figure 2. (A) Representative cryo-EM image and (B) 2D-class averages of vitrified mTMEM16F in a Ca^2+^-bound state in nanodiscs. (C) Angular distribution plot of particles included in the final C2-symmetric 3D reconstruction. The number of particles with their respective orientation is represented by length and color of the cylinders. (D) Image processing workflow. (E) FSC plot used for resolution estimation and model validation. The gold-standard FSC plot between two separately refined half-maps is shown in blue and indicates a final resolution of 3.5 Å. FSC validation curves for FSC_sum_, FSC_work_ and FSC_free_, as described in Methods, are shown as light blue, dark grey and light grey, respectively. A thumbnail of the mask used for FSC calculation overlaid on the atomic model is shown in the upper right corner and threshold used for FSC_sum_ of 0.5 and for FSC, FSC_work_ and FSC_free_ of 0.143 are shown as dashed lines. (F) Anisotropy estimation plot of the final map. The global FSC curve is represented in yellow. The directional FSCs along the x, y and z axis are shown in blue, green and red, respectively, and indicate a dataset that is to some extent anisotropic. (G) Final reconstruction map colored by local resolution as estimated by Relion, indicating regions of higher resolution. (H) Representation of the map including the nanodisc (low-pass filtered to 6.5 Å), reveals no noticeable distortion of the nanodisc (light blue). (I) Analysis of structural heterogeneity by subjecting the final dataset to another round of 3D classification, reveals alternative conformations of α3 and α4. Shown are superpositions of obtained 3D classes (grey) on the final cryo-EM map (light blue).

**Figure S5.**
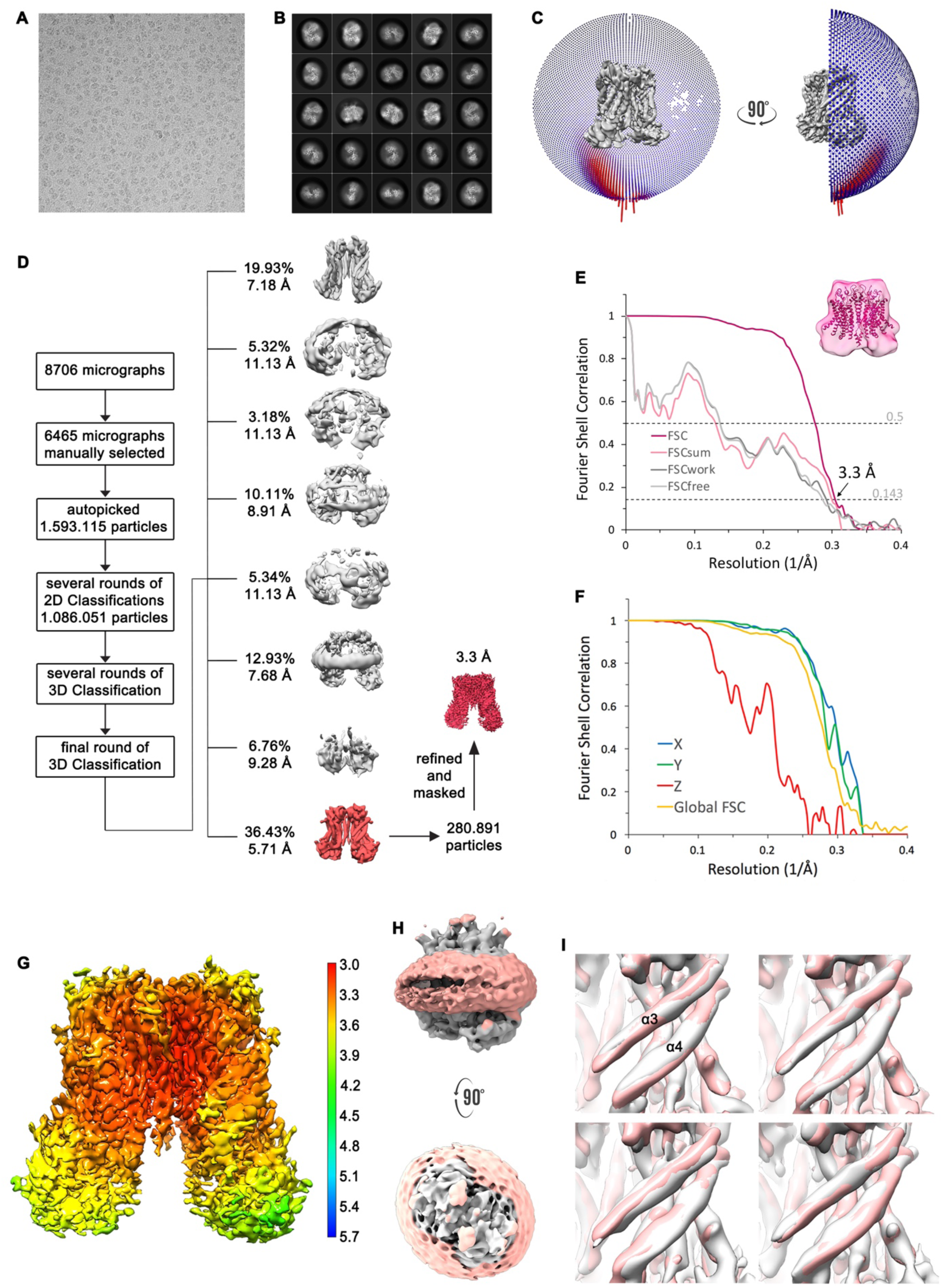
Structure Determination of TMEM16F in absence of Ca^2+^ in nanodiscs, Related to Figure 2. (A) Representative cryo-EM image and (B) 2D-class averages of vitrified mTMEM16F in a Ca^2+^-free state in nanodiscs. (C) Angular distribution plot of particles included in the final C2-symmetric 3D reconstruction. The number of particles with their respective orientation is represented by length and color of the cylinders. (D) Image processing workflow. (E) FSC plot used for resolution estimation and model validation. The gold-standard FSC plot between two separately refined half-maps is shown in red and indicates a final resolution of 3.3 Å. FSC validation curves for FSC_sum_, FSC_work_ and FSC_free_, as described in Methods, are shown as magenta, dark grey and light grey, respectively. A thumbnail of the mask used for FSC calculation overlaid on the atomic model is shown in the upper right corner and threshold used for FSC_sum_ of 0.5 and for FSC, FSC_work_ and FSC_free_ of 0.143 are shown as dashed lines. (F) Anisotropy estimation plot of the final map. The global FSC curve is represented in yellow. The directional FSCs along the x, y and z axis are shown in blue, green and red, respectively, reveal a strongly anisotropic dataset, which compromises a detailed interpretation of the map. (G) Final reconstruction map colored by local resolution as estimated by Relion. (H) Representation of the map including the nanodisc (low-pass filtered to 6.5 Å), reveals no noticeable distortion of the nanodisc (light red). (I) Analysis of structural heterogeneity by subjecting the final dataset to another round of 3D classification, reveals no alternative conformations of α3 and α4. Shown are superpositions of obtained 3D classes (grey) on the final cryo-EM map (light red).

**Figure S6.**
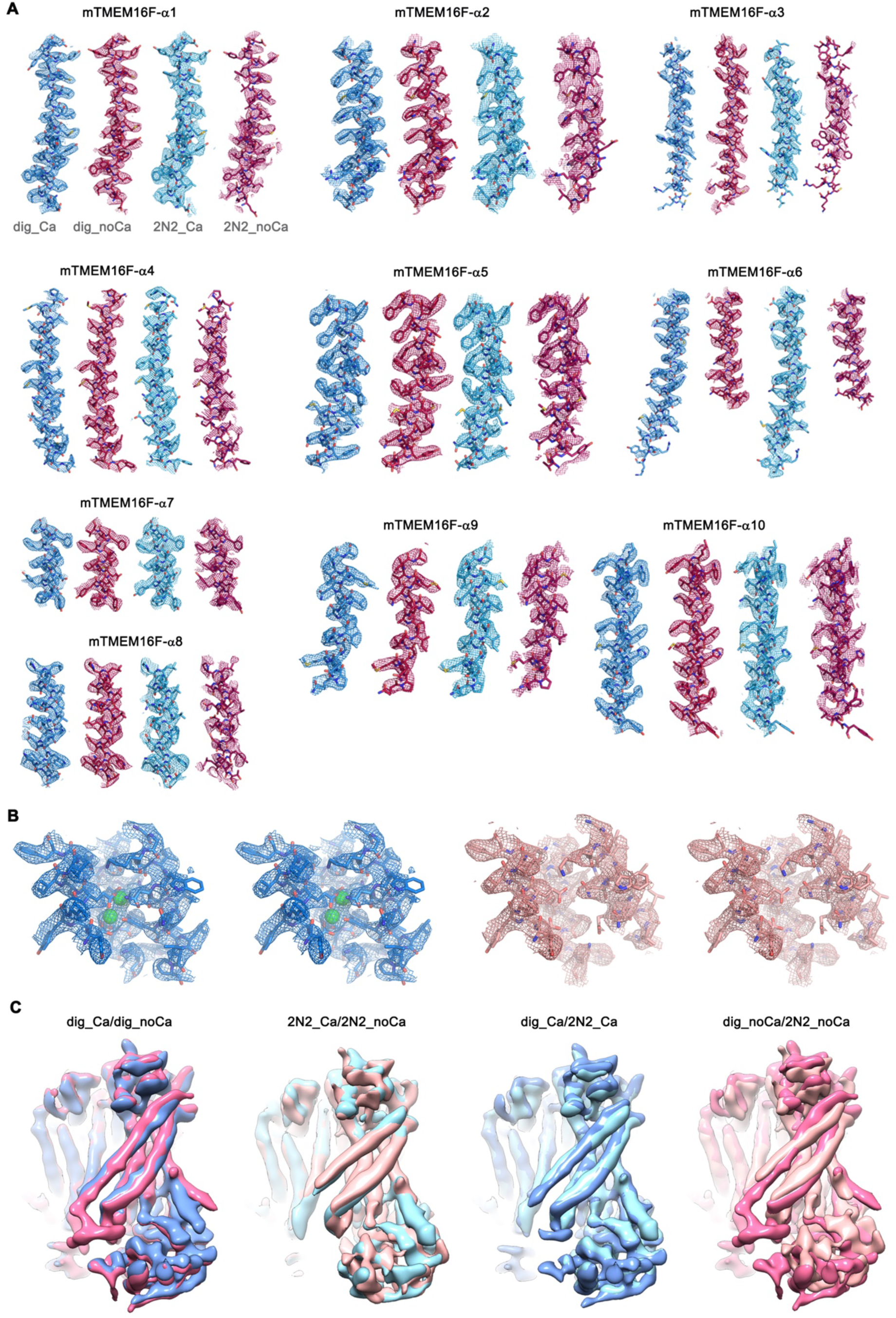
Cryo-EM Density Maps, Related to Figure 2. (A) Sections of the cryo-EM density of all four TMEM16F maps superimposed on the respective refined models. Models are shown as sticks and structural elements are labelled. TMEM16F Ca^2+^-bound in digitonin is coloured in blue, TMEM16F Ca^2+^-free in digitonin in red, TMEM16F Ca^2+^-bound in nanodiscs in light blue and TMEM16F Ca^2+^-free in nanodiscs in magenta. Maps were sharpened with a b-factor of –94, –139, –121 and –131 Å^2^ and contoured at 4.7 σ, 4.7 σ, 6.5 σ and 6.5 σ, respectively. (B) Stereo-view of the calcium-ion binding site as depicted in Figure 5B and 5C, B and C for the TMEM16F Ca^2+^-bound state in digitonin (left panel, blue) and the TMEM16F Ca^2+^-free state in digitonin (right panel, light brown). The respective cryo-EM density is shown as mesh, all residues are shown as sticks and Ca^2+^ as green spheres. Maps were sharpened with a b-factor of –94 and –139 Å^2^, respectively, and both contoured at 5 σ. (C) Superpositions of the four final cryo-EM maps. TMEM16F in presence of calcium is shown in dark blue in digitonin and in light blue when reconstituted in nanodiscs. TMEM16F in absence of calcium is shown in magenta in digitonin and in light magenta when reconstituted in nanodiscs. For a better comparison all maps were low-pass filtered to 6.5 Å. The conformational differences observed for α3 and α4 between the datasets obtained in digitonin and nanodiscs, might hint towards a lipid-dependent conformational transition.

**Figure S7.**
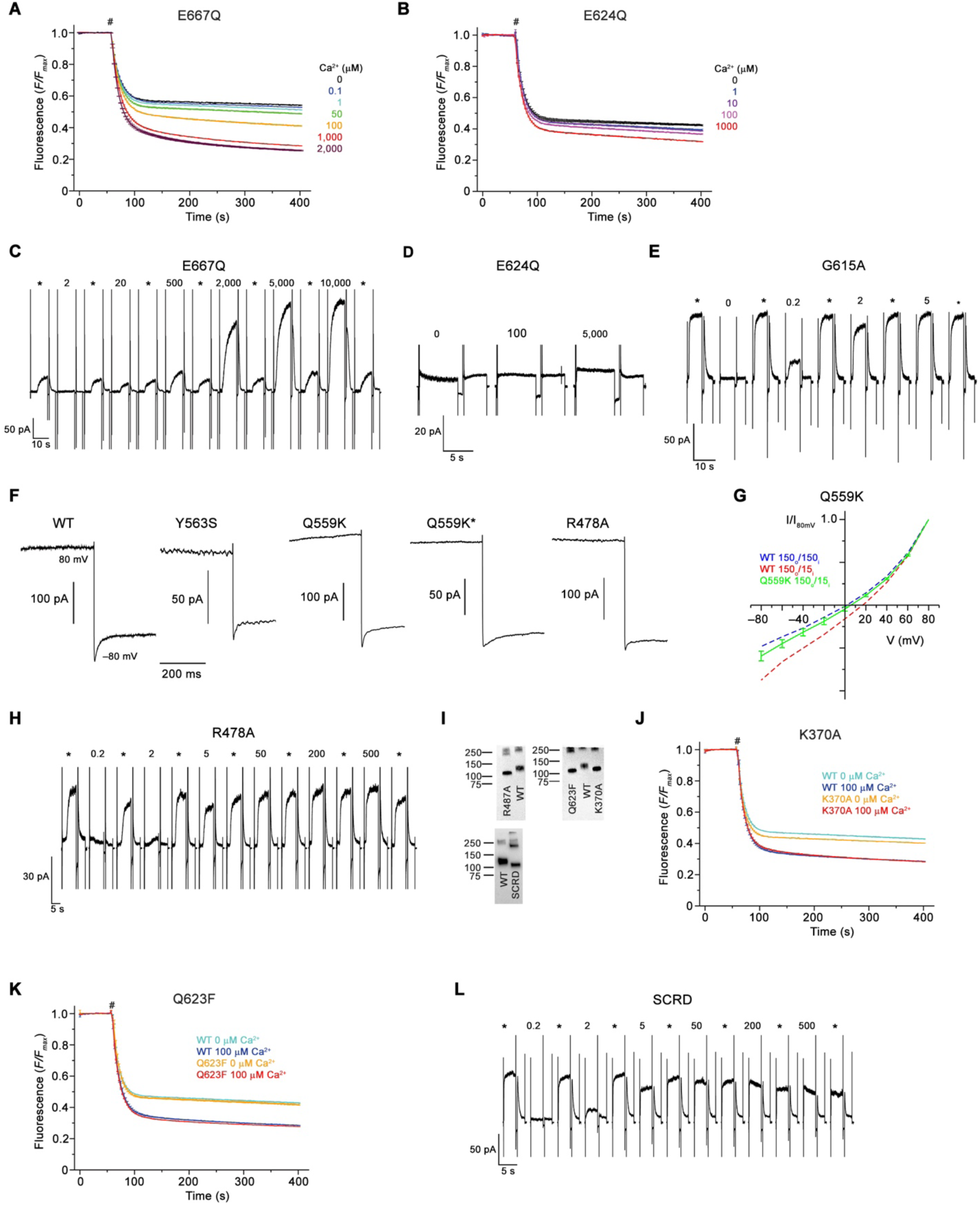
Electrophysiology and lipid scrambling data of mutants, Related to Figure 6. (A and B) Ca^2+^-dependence of scrambling activity in proteoliposomes containing the Ca^2+^-binding site mutants E667Q (A) and E624Q (B) at indicated ligand concentrations. (C, D and E) Representative current response of the TMEM16F mutants E667Q (C), E624Q (D), and G615A (E) at different Ca^2+^ concentrations. (F) Representative current traces of WT and mutants used for calculation of the rectification index (I_80mV_/I–_80mV_) of instantaneous currents (50 µs after voltage change) and at steady state. Q559K* corresponds to a subset of samples of the mutant with lower steady state rectification. (G) Normalized I-V relationships of the mutant Q559K in a 10-fold gradient of NaCl (green). The osmolarity of low NaCl solution was adjusted by addition of NMDG-sulfate. The shift in the reversal potential in asymmetric conditions towards 0 indicates a decreased cation over anion selectivity. WT at symmetric and asymmetric concentrations is indicated as dashed line for comparison. Data shows average of 5 biological replicates, errors are s.e.m.. (H) Representative current response of the TMEM16F mutant R478A at different Ca^2+^ concentrations. (I) Western blot analysis of the protein content of proteoliposomes containing different TMEM16F constructs after extraction with dodecyl-β-D-maltopyranoside. WT reconstituted in the same batch of liposomes as respective mutants was used as reference and is displayed on the same blot for comparison. Proteoliposomes of mutants show similar protein levels as respective WT controls and thus allow for a direct comparison of respective activities. (J and K) Scrambling activities of the mutant K370A (J) and Q623F (K) at indicated Ca^2+^ concentrations. (L) Representative current response of the construct TMEM16F^SCRD^ (SCRD) at different Ca^2+^ concentrations. (J and K) WT reconstituted in a separate set of liposomes from the same batch is shown as comparison. (C to E), (H and L) Data were recorded in excised inside-out patches. Numbers indicate Ca^2+^ concentrations (µM), * indicates reference pulses used for rundown correction recorded at 200 µM Ca^2+^ for E667Q and 20 µM Ca^2+^ for all other mutants. (A, B, J and K,) Traces depict fluorescence decrease of tail-labeled NBD-PE lipids after addition of dithionite (#). Data show averages of three technical replicates from the same reconstitution.

**Table S1.** Cryo-EM data collection, refinement and validation statistics, Related to Figure 2.

|  | mTMEM16F<br>Digitonin, +Ca<sup>2+</sup><br>(EMDB ...,<br>PDB ...) | mTMEM16F<br>Digitonin, -Ca<sup>2+</sup><br>(EMDB ...,<br>PDB ...) | mTMEM16F<br>2N2, +Ca<sup>2+</sup><br>(EMDB ...,<br>PDB ...) | mTMEM16F<br>2N2, -Ca<sup>2+</sup><br>(EMDB ...,<br>PDB ...) |
| --- | --- | --- | --- | --- |
| **Data collection and processing** |  |  |  |  |
| Microscope | Talos Arctica | Talos Arctica | Talos Arctica | Talos Arctica |
| Camera | K2 Summit + GIF | K2 Summit + GIF | K2 Summit + GIF | K2 Summit + GIF |
| Magnification | 49,407 | 49,407 | 49,407 | 49,407 |
| Voltage (kV) | 200 | 200 | 200 | 200 |
| Exposure time frame/total (s) | 0.15/9 | 0.15/9 | 0.15/9 | 0.15/9 |
| Number of frames per image | 60 | 60 | 60 | 60 |
| Electron exposure (e-/Å<sup>2</sup>) | 52 | 52 | 52 | 52 |
| Defocus range (μm) | -0.5 to -2.0 | -0.5 to -2.0 | -0.5 to -2.0 | -0.5 to -2.0 |
| Pixel size (Å) | 1.012 | 1.012 | 1.012 | 1.012 |
| Box size (pixels) | 220 | 220 | 256 | 220 |
| Symmetry imposed | C2 | C2 | C2 | C2 |
| Initial particle images (no.) | 1,348,247 | 1,314,676 | 1,019,012 | 1,593,115 |
| Final particle images (no.) | 219,302 | 194,284 | 186,487 | 280,891 |
| Map resolution (Å) | 3.18 | 3.64 | 3.54 | 3.27 |
| 0.143 FSC threshold |  |  |  |  |
| Map resolution range (Å) | 2.9-4.5 | 3.2-5 | 3.3-5 | 3.0-4.2 |
| **Refinement** |  |  |  |  |
| Initial model used | PDB 5OYB | PDB 5OYB | PDB 5OYB | PDB 5OYB |
| Model resolution (Å) | 3.3 | 3.8 | 4.0 | 7.0 |
| FSC threshold |  |  |  |  |
| Model resolution range (Å) | 80-3.2 | 80-3.6 | 80-3.5 | 80-3.3 |
| Map sharpening *B* factor (Å<sup>2</sup>) | -94 | -139 | -121 | -131 |
| Model composition |  |  |  |  |
| Nonhydrogen atoms | 10640 | 10566 | 10500 | 7810 |
| Protein residues | 1348 | 1336 | 1330 | 954 |
| Ligands | 4 | - | 4 | - |
| *B* factors (Å<sup>2</sup>) |  |  |  |  |
| Protein | 32.7 | 62.5 | 86.7 | 59.8 |
| Ligand | 19.8 | - | 67.9 | - |
| R.m.s. deviations |  |  |  |  |
| Bond lengths (Å) | 0.008 | 0.008 | 0.008 | 0.012 |
| Bond angles (°) | 1.37 | 1.146 | 1.091 | 1.384 |
| Validation |  |  |  |  |
| MolProbity score | 1.81 | 1.73 | 1.78 | 1.84 |
| Clashscore | 5.42 | 5.5 | 5.2 | 10.08 |
| Poor rotamers (%) | 0.37 | 0.75 | 0.38 | 0.71 |
| Ramachandran plot |  |  |  |  |
| Favored (%) | 90.93 | 93.4 | 91.42 | 95.48 |
| Allowed (%) | 8.92 | 6.6 | 8.58 | 4.52 |
| Disallowed (%) | 0.15 | 0 | 0 | 0 |

**Movie S1. Structure of TMEM16F in complex with Ca^2+^, Related to Figure 2**

Shown is the cryo-EM density map of TMEM16F obtained in complex with Ca^2+^ in digitonin superimposed on the refined structure. For clarity, only a single subunit is shown. The cryo-EM map is depicted as blue surface, the model as blue sticks, and calcium ions as green spheres.

**Movie S2. Structure of TMEM16F in absence of Ca^2+^, Related to Figure 2**

Shown is the cryo-EM density map of the TMEM16F obtained in a Ca^2+^-free state in digitonin superimposed on the refined structure. For clarity, only a single subunit is shown. The cryo-EM map is depicted as magenta surface and the model as red sticks.

